# Surface tension-driven sorting of human perilipins on lipid droplets

**DOI:** 10.1101/2024.02.12.579497

**Authors:** Ana Rita Dias Araujo, Abdoul Akim Bello, Joëlle Bigay, Céline Franckhauser, Romain Gautier, Julie Cazareth, David Kovacs, Frédéric Brau, Nicolas Fuggetta, Alenka Copic, Bruno Antonny

## Abstract

Perilipins (PLINs), the most abundant proteins on lipid droplets (LDs), display similar domain organization including amphipathic helices (AH). However, the five human PLINs bind different LDs suggesting different modes of interaction. We established a minimal system whereby artificial LDs covered with defined polar lipids were transiently deformed to promote surface tension. Binding of purified PLIN3 and PLIN4 AH was dependent on tension, even with polar lipids favoring packing defects, and showed an inverse correlation between protein and phospholipid densities on LDs. In contrast, PLIN1 bound readily to LDs fully covered by phospholipids; PLIN2 showed an intermediate behavior. In human adipocytes, PLIN3/4 were found in a soluble pool and relocated to LDs upon stimulation of triglyceride synthesis, whereas PLIN1 and PLIN2 localized to pre-existing LDs, consistent with the huge difference in LD avidity observed *in vitro*. We conclude that the PLIN repertoire is adapted to handling LDs with different surface properties.

**Significance statement:** Lipid droplets (LDs) are highly dynamic organelles, whose size and surface properties vary during their life-time and also differ between different tissues. Here, we analyze the mode of binding of human perilipins (PLINs), the most abundant LD proteins, to LDs. We have developed a new reconstitution method, which shows that the purified PLIN family members have very different affinities for LDs, which might explain how they handle LDs of different dynamics in the cell.

## Introduction

Lipid droplets (LDs) differ markedly from other cellular organelles in that they are not surrounded by a phospholipid bilayer but by a monolayer, which isolates the LD core made of triolein and other neutral lipids from the cytosol (1, 2). The LD monolayer displays a phospholipid composition similar to that of the ER from which LDs form (3, 4). Therefore, preferential targeting of some proteins to LDs might not result from their recognition of specific lipids but rather from the unique biophysical properties of LDs (1, 2, 5). Notably, the phospholipid density of the LD monolayer should vary according to the cellular metabolic state (e.g. neutral lipid synthesis *vs* lipolysis), resulting in differences in LD surface tension. Molecular dynamic simulations suggest that at high surface tension, triolein molecules interdigitate with the phospholipid acyl chains or even participate in the monolayer by adopting an ordered conformation, creating a lipid surface very different from that of a bilayer (5, 6). Furthermore, the size and thus curvature of LDs varies over several orders of magnitude between different cells, and also within the same cell, for example in adipocytes during differentiation. In the case of bilayer-bound organelles, curvature acts as a targeting signal for proteins that detect lipid packing defects present in the bilayer (7). Similar packing defects can be induced by changes in lipid composition, for example by increased concentration of lipids with small headgroups such as phosphatidylethanolamine or diacylglycerol (8). Whether the packing defects present on small LDs, on LDs under tension or on LDs undergoing changes in lipid composition are similar to those present in lipid bilayers is elusive and difficult to tackle experimentally (5, 6, 9). Overall, targeting of proteins to LDs might be governed by rules different from those observed for other organelles, requiring models to recapitulate the unique LD features.

PLINs are highly abundant LD proteins with no enzymatic activity. Due to their high abundance and ability to stabilize LDs, PLINs are often referred to as LD coats (10–12). In humans, the PLIN family contains 5 members (PLIN1-5), which show notable differences in their tissue and subcellular distribution, mode of regulation, and protein interactions (12–14). PLINs contain three main regions: (i) an N-terminal PAT domain, the structure of which is not known but which might contain three helices (15), (ii) a central disordered region, which has been shown to target lipid droplets by providing 3-11 amphipathic helices (AH) that bind to the LD interface (16–20), and (iii) a C-terminal domain, whose structure has been solved in the case of PLIN3 as forming a 4-helix bundle (21). However, PLINs show variations around this canonical scheme. For instance, the 4-helix bundle region of PLIN1 is not well conserved, containing hydrophobic segments that are required for LD targeting in cells(22), and is followed by a C-terminal extension critical for lipolysis control in white adipocytes(23, 24). PLIN4 does not contain a PAT region, whereas its central AH is dramatically extended, containing 29 x 33-mer repeats (19, 20, 25). PLIN5 contains a C-terminal extension that allows it to tether LDs to mitochondria (26). Although progress has been made towards deciphering the various PLIN regions, the presence of intrinsically disordered regions and extended lipid-interacting surfaces, as well as the low solubility of some PLINs have hampered in-depth structural characterization. Therefore, our understanding of the mechanisms by which PLINs interact with the surface of LDs remains fragmental.

One long-standing question is why perilipins appear heterogeneously distributed on LDs. In 2D cell cultures of adipocytes, there is a gradient of LD coverage by PLINs. PLIN1 outlines large and centrally localized LDs, PLIN2 is found at the surface of intermediate LDs, whereas PLIN3 and PLIN4 mark small and peripheral LDs (Wolins et al., 2006; 2005; 2003). This uneven distribution might result from differences in protein turnover during cell differentiation and/or from intrinsic differences in the physicochemical properties of PLINs. These include the amino-acid composition of their AH regions, the presence of additional hydrophobic regions, or the ability of the PAT region or the 4-helix bundle to undergo conformational changes (14, 15, 20, 27).

Directly addressing the physicochemical properties of PLINs has been difficult. Only PLIN2/3 have been purified in their full-length form (28, 29). Furthermore, artificial LDs for biochemical reconstitution are far from being mastered as well as artificial lipid bilayers, although significant progress has been made in recent years (2, 30–33). In this work, we set up a new LD mimetic system amenable to transient tension challenge and to analytical assays including flow cytometry for quantification of protein binding. We show that purified full-length PLIN1/2/3 display strikingly different LD binding properties, consistent with their different LD targeting in cells.

## Results

### Rationale for LD preparation and protein binding protocols

In order to study the interaction of PLINs with LDs, we aimed to set up a robust LD mimetic protocol with which we could perform complementary assays including visualization by fluorescence microscopy, bulk biochemical assays, and flow cytometry analysis to gather quantitative data from all individual LDs in a mixture. However, manipulating LDs in vitro is challenging. First, the oil density (≈ 0.9 g/mL) makes LDs light objects that readily float in a microscopy chamber or in a test tube, which necessarily impedes homogenous contact with the bulk phase. Second, gentle addition of a protein to a pre-formed oil droplet covered with phospholipids might lead to a false negative (i.e., no binding) result because the phospholipid surface is too dense to accommodate the protein, whereas the protein may be well adapted to binding to LDs with a lower surface density. The latter consideration is important for PLINs, which are the most abundant LD-associated proteins and contain long (a hundred aa) or very long (a thousand aa) AH motifs (10, 11, 19, 34). The amount of phospholipids necessary to make stable preformed LDs might hinder the subsequent binding of such proteins with an extended interfacial surface.

To circumvent these difficulties, we established a new LD binding assay that allowed us to both control the LD density and apply a deformation step to transiently increase surface tension (**Figure 1A**). The LD core was made of a mixture of triolein (TG(18:1/18:1/18:1)) and of a heavy oil that is denser than water (brominated vegetable oil, BVO). By adjusting the ratio of the two oils, one can prepare artificial droplets with a defined density (33). We chose a density of 1.05 g/mL, which corresponded to a brominated oil:triolein ratio of 1:2. Thus, triolein dominated the oil phase but the droplets had a slightly higher density than the buffer, which enabled sedimentation of LDs in the optical chamber. Phospholipids were added at a phospholipid/oil ratio that gave stable LDs with a defined average diameter, which were further sized by an extrusion step (see methods). We chose an average LD diameter of 2 µm for flotation and flow cytometry experiments, and 10 µm for light microscopy experiments. This diameter range is similar to that of many cellular lipid droplets.

**Figure 1.**
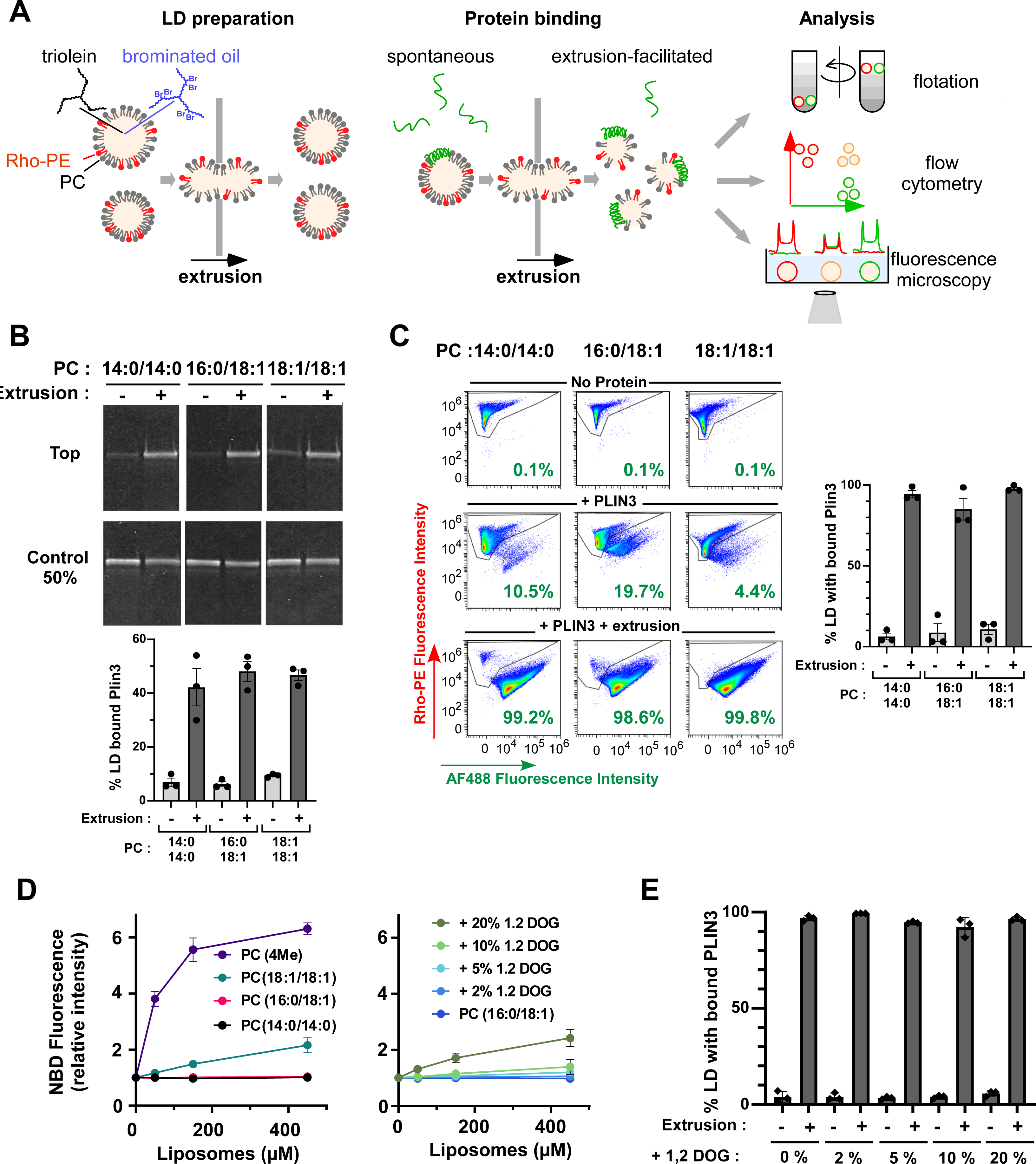
Binding of PLIN3 to artificial LDs requires surface tension. **A.** Rationale of the LD binding assay. Artificial PC-covered lipid droplets containing a mix of heavy (BVO) and normal (TG) oil were sized by a first extrusion step. Thereafter, a protein of interest was gently added. Half of the sample was processed directly for analysis. The other half was submitted to a second extrusion step to transiently increase LD surface tension. Three complementary methods were used for analysis: flotation to determine the fraction of LD-bound protein; flow cytometry to determine the fraction of LDs with bound protein; fluorescence microscopy to determine the surface density of proteins *vs* phospholipids. **B.** Flotation analysis. 1.4 µM PLIN3 (of which 5 % was labeled with AF488) was mixed with PC-covered lipid droplets (volume fraction of oil = 0.75%; calculated diameter = 2 µm; concentration of phospholipid [PL] = 62.5 µM). When indicated, the mixture was further extruded through 1 µm polycarbonate filters. After flotation, the top fraction was analyzed by SDS-PAGE and direct AF488 fluorescence detection. Bar plot: quantification (mean ± SEM) of the fraction of LD-bound AF488 PLIN3 (n= 3). **C.** Flow cytometry analysis of PLIN3 + LD samples similar to that used in B. Bar plot: quantification (mean ± SEM) of the fraction of AF488 PLIN3-positive LDs (n = 3). **D**. Liposome-binding assays showing the fluorescence (mean ± SEM, N = 3) of NBD PLIN3 in the presence of increasing amounts of liposomes. Left plot: experiments performed with PC liposomes of defined acyl chain composition. Right plot: experiments performed with PC(16:0/18:1) liposomes containing increasing amounts of 1,2 Dioleoylglycerol (DOG). DOG slightly favors the binding of NBD PLIN3 to PC liposomes. **E**. DOG has no effect on the binding of AF488 PLIN3 to LDs, as determined by flow cytometry under conditions similar to B and C (mean ± SEM, N = 3).

We compared protein binding to these LDs under two conditions: when proteins were gently added to LDs, or when we applied a second extrusion step to the LD-protein mixture (**Figure 1A**). In this second step, we forced the LD-protein mix to pass through a polycarbonate filter of a smaller pore size than the average diameter of the initial LD suspension (e.g. 1 µm *vs* 2 µm or 8 µm vs 10 µm). This should create a transient tension at the LD interface by deforming the initially spherical LDs. The resulting increase in the surface to volume ratio might facilitate protein insertion.

### PLIN3 binding to LD is hypersensitive to surface tension

We first performed experiments with full-length PLIN3 and with phosphatidylcholine (PC)-covered LDs (**Figure 1B**). PLIN3 is intrinsically soluble and its purification does not require any detergent or chaotropic agent (15, 29, 34). PC is the most abundant phospholipid found on LDs in cells (3, 4). To facilitate the analysis, PLIN3 was labelled with AlexaFluor488 (AF488) and used at 5 - 20 mol % compared to unlabeled PLIN3, whereas the phospholipids included a fraction of 16:0 lissamine rhodamine B phosphatidylethanolamine (RhoPE) as a fluorescent tracer. Because acyl chain unsaturation can strongly affect the interaction of AH-containing proteins with lipid membranes (35), we used PC species of increasing level of unsaturation, namely PC(14:0/14:0) (fully saturated), PC(16:0/18:1) (saturated-monounsaturated), and PC(18:1/18:1) (di-monounsaturated).

We incubated purified PLIN3 for 10 min with LDs, which were subsequently isolated by flotation on sucrose cushions. Under these conditions, only a small fraction of PLIN3 was recovered in the top fraction containing the LDs. We observed slightly more binding to PC(18:1/18:1)-covered LDs (≈ 10%) than to PC(14:0/14:0)- or PC(16:0/18:1)-covered LDs (≈ 5%) (**Figure 1B**). In contrast, the level of PLIN3 recovery in the LD fraction dramatically increased upon extrusion, reaching values close to 50% regardless of the PC species (**Figure 1B**). This increase was observed for both AF488 PLIN3, as directly visualized by SDS-PAGE before staining, and for total PLIN3, as detected after total protein staining (**Fig. S1A**), suggesting that fluorescent labelling did not modify the affinity of the protein for LDs. High performance thin layer chromatography (HPTLC) (**Fig. S1B**) and SDS-PAGE (**Figure 1B**) showed that the majority (≈ 75%) of lipids and of protein was recovered after extrusion.

We next conducted flow cytometry measurements on similar LD-protein mixtures. Plotting the side (SSC) *vs* forward light scattering signals (FSC) of the various LD-protein suspensions showed an S shape, which is characteristic of well-defined lipid emulsions (**Fig. S1C**) (36). Analyzing the RhoPE channel to follow droplets and the AF488 channel for PLIN3 before extrusion showed that only a small fraction of the droplets exhibited the protein signal (≈ 10%). After extrusion, the large majority of LDs (≈ 90%) displayed the two signals, confirming the dramatic effect of extrusion on PLIN3 recruitment to the LD surface (**Figure 1C**).

Next, we performed several complementary experiments. First, we used either larger LDs or omitted brominated oil in the LD formulation. These modifications did not change the behavior of PLIN3, which remained strongly dependent on extrusion for its binding to LDs (**Fig. S1D**), suggesting that the presence of brominated oil did not modify the surface properties of the LDs. Second, we varied the ratio between labelled and unlabelled PLIN3 and observed similar results, confirming that PLIN3 labelling did not modify LD binding properties (**Fig. S1E**). Third, we compared various fragments of PLIN3. We observed that the AH region of PLIN3 (aa 114-204) and a construct encompassing both the PAT domain and the AH region of PLIN3 (aa 1-205) showed similar LD binding properties as full-length PLIN3, including a strong dependency on extrusion (**Fig. S1F**). These results matched well with previous observations that the AH region of PLIN3 is the main determinant for LD binding (17, 18, 20, 27). We also performed LD binding experiments with a fragment of PLIN4 that encompasses a large part of its giant AH region (aa 246-905, PLIN4 AH-20mer) (19, 20). This fragment behaved similarly to PLIN3 in the LD-binding assay, showing almost no spontaneous binding to PC-covered LDs and strong binding after extrusion (**Fig. S1F**).

Finally, we tested the effect of 1,2 dioleoyl-glycerol (DOG) on PLIN3 binding properties. Diacylglycerols are intermediates in LD metabolism and have been shown to promote the binding of PLIN3 to lipid bilayers (15, 37–39). To quantify the effect of DOG on PLIN3 binding to bilayers, we used the membrane environment-sensitive fluorescent probe NBD (19). We incubated NBD-labelled PLIN3 with liposomes of defined lipid composition and used NBD fluorescence intensity as an index of protein binding (**Figure 1D**). NBD PLIN3 did not bind to PC(14:0/14:0) or PC(16:0/18:1) liposomes, showed modest binding to PC(18:1/18:1) liposomes, whereas strong binding was observed on diphytanoyl (4Me) PC liposomes. The branched acyl chains of PC(4Me) promote large lipid packing defects in membranes, which facilitate the binding of AHs, including the long AH of PLIN4 (19, 40). Increasing the amount of DOG at the expense of PC(16:0/18:1) led to a modest increase in PLIN3 binding to liposomes, which became significant only at DOG levels > 10 mol%. We then asked whether DOG could alleviate the need for tension in the case of PLIN3 binding to LDs. Surprisingly, increasing the amount DOG in the LD formulation had no effect on the binding of PLIN3 to LDs, which remained highly dependent on extrusion (**Figure 1E**). We conclude that PLIN3 does not bind spontaneously to LDs even when they are surrounded by a DOG-enriched lipid monolayer, whereas it binds avidly to LDs under tension.

### Systematic LD-binding measurements indicate a very large affinity range among PLINs

We next wanted to evaluate the binding of other PLINs to LDs. However, they were more challenging to express and purify than PLIN3. PLIN1, PLIN2 and PLIN5 were insoluble in standard buffers, and full-length PLIN4 could not be expressed in *E. coli* due to its size. To overcome these difficulties, we set up different expression and purification procedures (**Fig. S2A**). PLIN1 was expressed at low temperature in *E. coli* arctic cells, and the solubility of PLIN1 and PLIN2 was improved by the addition of urea (**Fig. S2A**). For PLIN4, we used constructs that encompass most of its AH region and which can be readily purified (19, 20). PLIN5 was not included in the analysis.

We compared the binding of full-length PLIN1, PLIN2 and PLIN3 to PC-covered LDs using the flow cytometry approach (**Figure 2A**). PLIN1 and PLIN2 were tested in the presence of urea at a concentration (2M) that was sufficient to keep the two proteins soluble (**Fig. S2B**). Control experiments showed that binding of PLIN3 to LDs was not affected by the presence of 2M urea and remained highly promoted by extrusion (**Fig. S2C**). In addition, 2M urea did not modify the pattern and kinetics of limited proteolysis of PLIN1, PLIN2 and PLIN3 by subtilisin. At both 0 and 2M urea, PLIN2 and PLIN3 showed a proteolysis-resistant C-terminal domain, corresponding to the C-terminal 4-helix bundle, whereas PLIN1 was susceptible to proteolysis along its entire length/sequence (**Fig. S2D**). These data suggest that the overall domain organization of the three perilipins is not affected by 2M urea.

**Figure 2.**
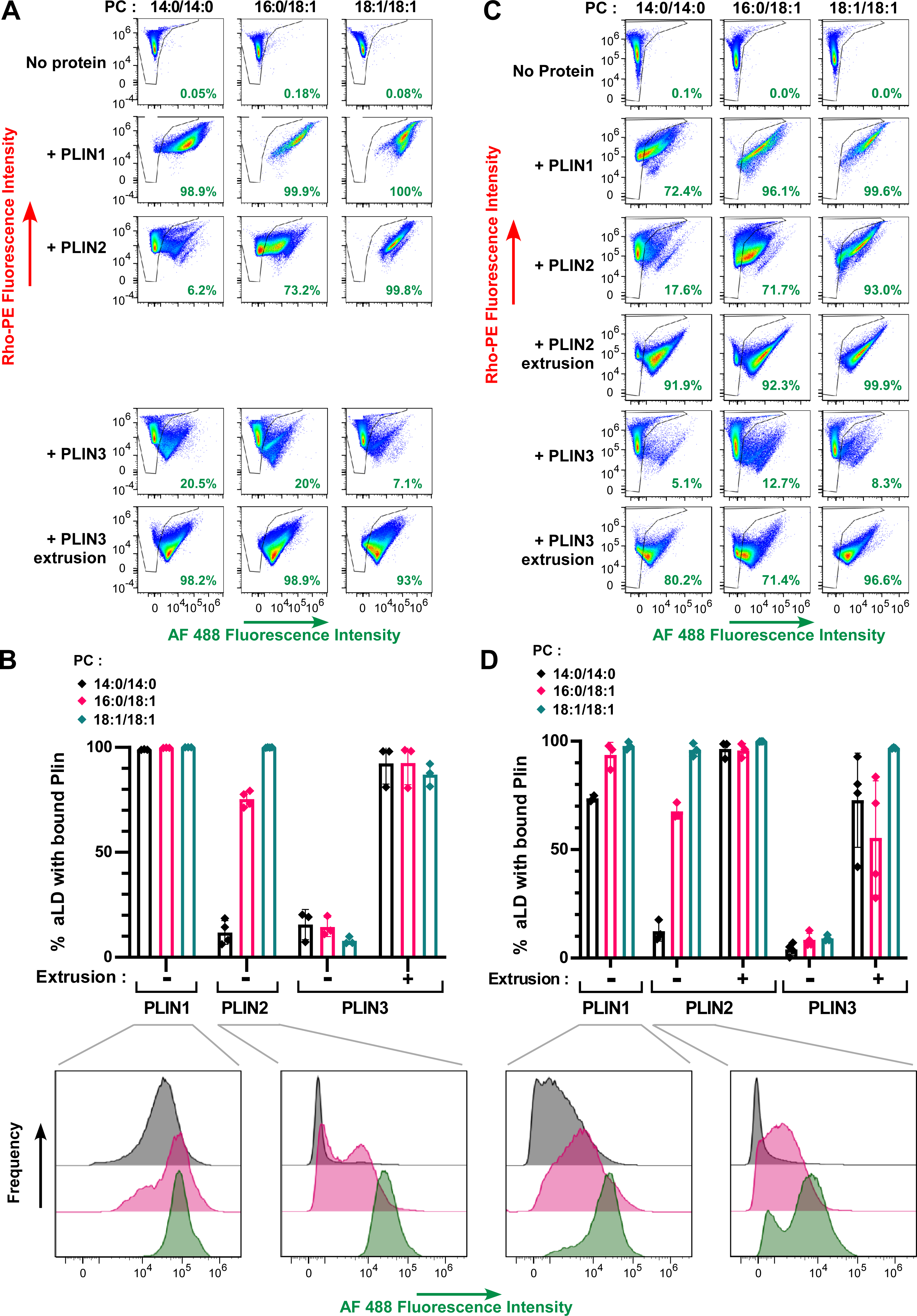
Binding of full-length PLIN1 and PLIN2 to artificial LDs. Flow cytometry analysis of PLIN1, PLIN2 or PLIN3 in the presence of PC-covered LDs (2 µm). When indicated, protein binding was analyzed before and after extrusion. The protein concentration was 1.4 µM (**A, B**) and 0.35 µM (**C, D**). Bar plots in B and D show quantification (mean ± SEM) of the fraction of AF488 PLIN positive LDs from experiments similar to that shown in A and C, respectively (n = 3). For some conditions, the AF488 intensity distribution is shown to illustrate differences in PLIN density on the PLIN positive LDs.

**Figure 2A** shows LD binding experiments with full-length PLIN1, PLIN2 or PLIN3 as analyzed by flow cytometry. PLIN1 bound spontaneously to LDs regardless of the PC species (14:0/14:0, 16:0/18:1 or 18:1/18:1) at the surface. However, the AF488 signal was less intense on the PC(14:0/14:0)-covered LDs, suggesting lower protein density (**Figure 2B**). For PLIN2, we observed a large effect of PC unsaturation, with the fraction of PLIN2-positive LDs increasing from 10% with PC(14:0/14:0)-covered LDs up to 100% with PC(18:1/18:1)-covered LDs (**Figure 2A, B**). Because of the low spontaneous binding of PLIN2 to LDs covered with saturated PC, we aimed to test the effect of extrusion. However, we noticed that a part of the LD-PLIN1 and LD-PLIN2 particles might have aggregated because the light scattering diagrams did not show a well-defined S-shape (**Figure S2E**). Excess protein in the case of PLIN1 and PLIN2 might be less tolerated compared to PLIN3, which is very soluble. We thus repeated these experiments with a four-fold lower protein concentration and observed very similar results in terms of % of PLIN-positive LDs, although we observed a lower AF-488 signal suggesting less bound-protein, as expected (compare **Figure 2C, D** and **Figure 2A, B**). We then applied extrusion to the PLIN2-LD samples and observed that all LDs became PLIN2 positive (**Figure 2C, D**), suggesting that PLIN2 is also sensitive to LD surface tension but that this requirement can be bypassed when the PC monolayer is sufficiently unsaturated. Altogether, these experiments indicated that the three PLINs have very different affinities for PC-covered droplets, with the order PLIN3 << PLIN2 << PLIN1. We note that this order matches their calculated hydrophobicity (20, 27) and experimentally measured solubility in the presence or absence of urea (**Fig. S2B**).

### Immunofluorescence in human adipocyte cultures suggests that PLINS have different roles

Given the striking difference between PLINs in their sensitivity to extrusion for binding to artificial LDs, we wanted to compare their binding to LDs in cells. For this, we chose the human adipocytes hWA, which can be differentiated in cell culture dishes and which endogenously express human PLIN1-4 (41, 42). We applied an optimized differentiation protocol to these cells, which led to efficient differentiation of most cells in the culture dish over the course of 14 days, as shown by the progressive enlargement of LDs observed by transmission microscopy (**Figure S3A**) and the expression of the adipocyte marker hormone-sensitive lipase (HSL) (**Figure 3A**). We observed a large concomitant increase in the expression of adipocyte-specific PLIN1 and PLIN4, as shown by Western blot analysis using antibodies against endogenous proteins (**Figure 3A**) in accordance with previous observations (43). We were not successful in probing for PLIN3, which is expressed in non-differentiated cells and whose expression decreases during differentiation (44). PLIN2 could be detected already in non-differentiated cells and showed a modest increase during differentiation (**Figure 3A**), as reported.

**Figure 3.**
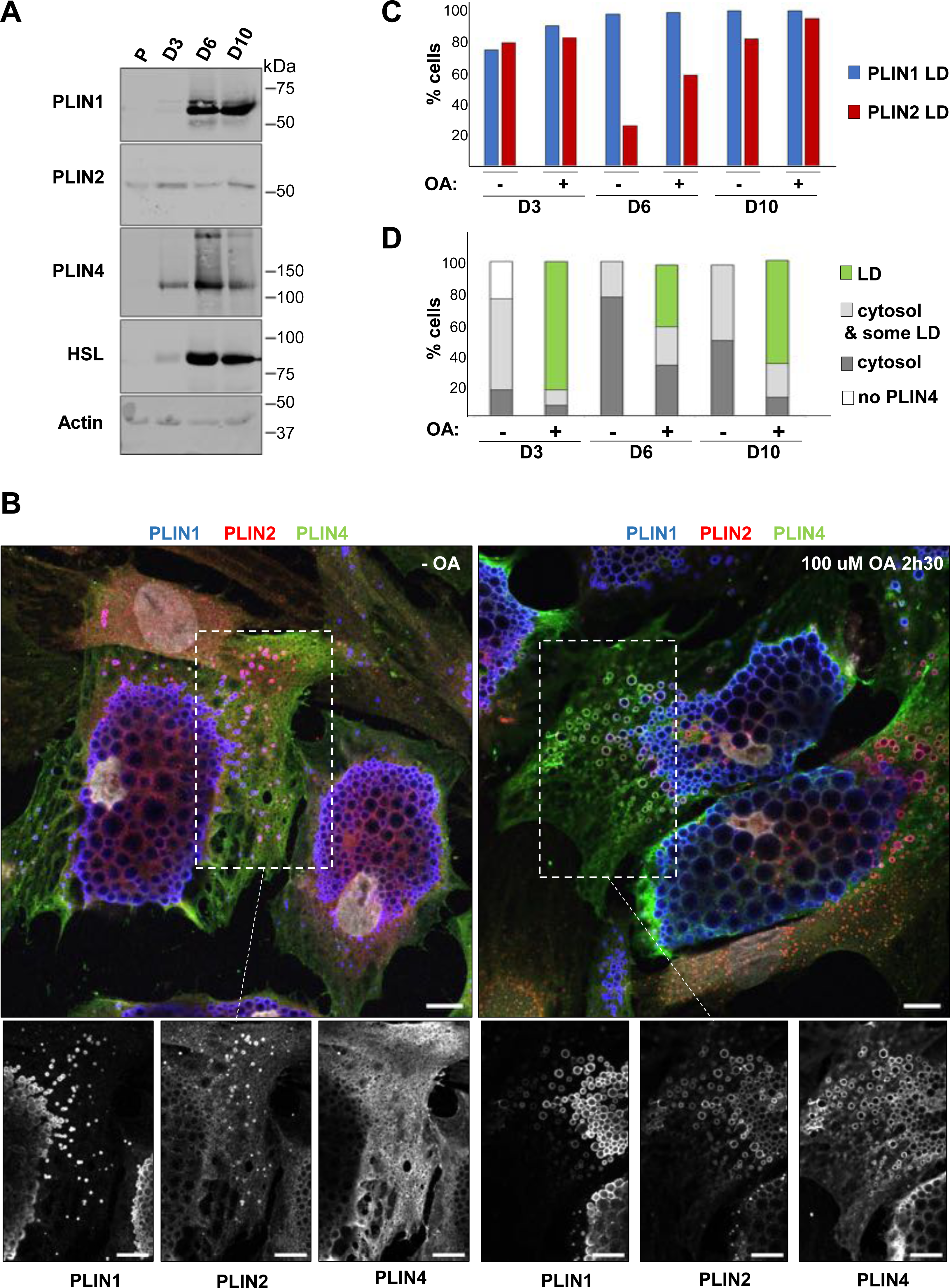
Analysis of perilipin expression and distribution in human adipocytes. **A.** Western blot analysis of adipocyte markers and PLINs during adipocyte differentiation. P: proliferation; D3, D6 and D11: day of differentiation. **B.** Representative z sections obtained by confocal fluorescence microscopy of endogenous PLIN1 (blue), PLIN2 (red) and PLIN4 (green) in human adipocytes at day 10 of differentiation after immunofluorescence with specific antibodies, with the nuclei stained with DAPI (white). Bottom panels show the three protein channels in the indicated area. The cells were either left in culture medium or fed with 100 µM oleic acid for 2.5 hours before observation. Scale bar: 10 µm. **C.** Quantification of the fraction of cells in which the PLIN1 and PLIN2 signals localized to LDs at D3, D6 and D10 of differentiation. **D.** Quantification showing the fraction of cells with PLIN4 signal as indicated under the same conditions as in B. Four categories were defined. Cells without detectable PLIN4; cells in which PLIN4 was entirely cytosolic; cells in which PLIN4 was mostly cytosolic but marked a few LDs; cells in which PLIN4 was LD-localized. N of cells quantified was 108 for D3 - OA, 104 for D3 + OA, 92 for D6 - OA, 92 for D6 + OA, 98 for D10 - OA and 93 for D10 + OA, from one of two representative experiments. See Figure S3B, C for representative immunofluorescence images from D3 and D6 of adipocyte differentiation.

We then analyzed the localization of different PLINs by immuno-fluorescence (IF) followed by 3D confocal imaging. For this analysis, we selected three different time-points of the differentiation protocol, early (day 3), intermediate (day 6), and late (day 10), correlating with the largest increase in LD size. We assessed the localization of PLIN1, PLIN2 and PLIN4 in the same cells under two conditions; in normal growth media or after the addition of 100 μM oleic acid (complexed with BSA; OA) for 2.5 h before fixation and IF analysis (**Figure 3B and Figure S3B-C**). Under all conditions, we observed PLIN1 on centrally localized LDs whose size progressively increased with differentiation, but did not show a large change after OA treatment. In striking contrast, PLIN4 displayed diffuse cytosolic signal in almost all non-treated cells at all stages of differentiation, with only faint staining of some peripheral LDs in a fraction of cells. Upon addition of OA, PLIN4 relocalized to peripheral LDs in a large fraction of cells (50% or more), with only a low number of cells retaining only cytosolic signal (**Figure 3C**). Importantly, this effect was observed both early and late in differentiation. There was some heterogeneity in the differentiating adipocyte population, and we observed that, on the same day, in the cells with less abundant or smaller PLIN1-labelled LDs, PLIN4 more frequently relocalized to small LDs after OA addition. PLIN2 displayed an intermediate behavior: it could be observed in the cytosol and on more peripheral/smaller LDs in the absence of OA; it decorated LDs more strongly after OA addition (**Figure 3B-C** and **Figure S3B-C**). We also analyzed the localization of PLIN3 at day 6 and observed behavior very similar to PLIN4, with cytosolic PLIN3 signal in the absence of OA, and an even higher LD localization after OA addition (**Figure S3D**).

These results are in agreement with previous observations in mouse differentiated 3T3-L1 adipocyte cells that showed relocalization of PLIN4 and PLIN3 to small peripheral LDs after OA addition, leading to the conclusion that these PLINs specifically localize to nascent LDs (43, 45). However, our analysis at different stages of adipocyte differentiation instead suggests that PLIN4 and PLIN3 function as buffers, localizing to newly-formed LDs after a burst in the production of triglycerides that is induced by OA addition, but they do not bind to LDs that form gradually during adipocyte differentiation.

### Visualization of PLIN3/4 binding to LDs reveals phospholipid exclusion

We wished to better understand the basis for the differences in PLIN recruitment to artificial or cellular LDs. From the flow cytometry analysis (see **Figure 1C**), we noticed that PLIN3 binding to LDs was accompanied by a large (5 to 10-fold) decrease in the RhoPE signal, suggesting that PLIN3 binding correlated with a reduction in LD size and/or phospholipid density. A similar trend was observed for PLIN2 (**Figure 2**). We thus determined the size of the LDs by dynamic light scattering measurements as well as by directly visualizing the LDs in a flow cytometry apparatus equipped with a camera (**Fig. S4A** and **B**). Whereas the mere addition of PLIN3 did not modify LD size, the combination of PLIN3 addition and LD extrusion led to a significant reduction in LD diameter. This observation suggests that, in addition to phospholipids, PLIN3 makes a significant contribution to the surface of LDs, hence enabling the formation of smaller LDs.

To directly visualize the protein and the phospholipid density on the artificial LDs, we used large PC(16:0/18:1)-covered LDs, which could be observed by light microscopy. **Figure 4A** compares typical microscopy fields of LDs alone, LD + PLIN3, and LD + PLIN3 after extrusion. In the absence of protein, all LDs were delimited by a similar phospholipid signal, as visualized in the RhoPE channel. Upon gentle addition of PLIN3, about one third of the droplets appeared covered by AF488 PLIN3, whereas the remaining LDs seemed devoid of it. Interestingly, the PLIN3-positive droplets exhibited a weak RhoPE signal as compared to the PLIN3-negative droplets. After extrusion, we could no longer distinguish different LD populations, but rather observed a continuum of LDs displaying various levels of AF488 and RhoPE fluorescence on their contour. To quantify the relative coverage of the LDs by PLIN3 and by phospholipids, we determined the red and green fluorescence profiles of hundreds of LDs and ranked them according to their coverage by AF488 PLIN3 (**Figure 4A**). The analysis confirmed the presence of two LD populations before extrusion. The first population (≈ 65% of LDs) displayed a phospholipid density similar to the initial LD preparation and a PLIN3 signal that did not exceed 10% of the signal of the protein in solution. The second droplet population (≈ 35%) displayed a clear PLIN3 signal, but with a lower RhoPE signal compared to the PLIN3-negative LDs or to the initial LD preparation. For this second population, the RhoPE signal and the AF488 signal correlated in an inverse manner; at maximal PLIN3 intensity, the RhoPE intensity dropped to ≈ 20% of the initial signal (**Figure 4B**). After extrusion, almost all droplets displayed AF488 PLIN3 whereas the RhoPE signal was 3.5 times lower than initially (**Figure 4B**). This analysis suggests that PLIN3 binding to LDs correlates with a strong reduction in phospholipid density.

**Figure 4.**
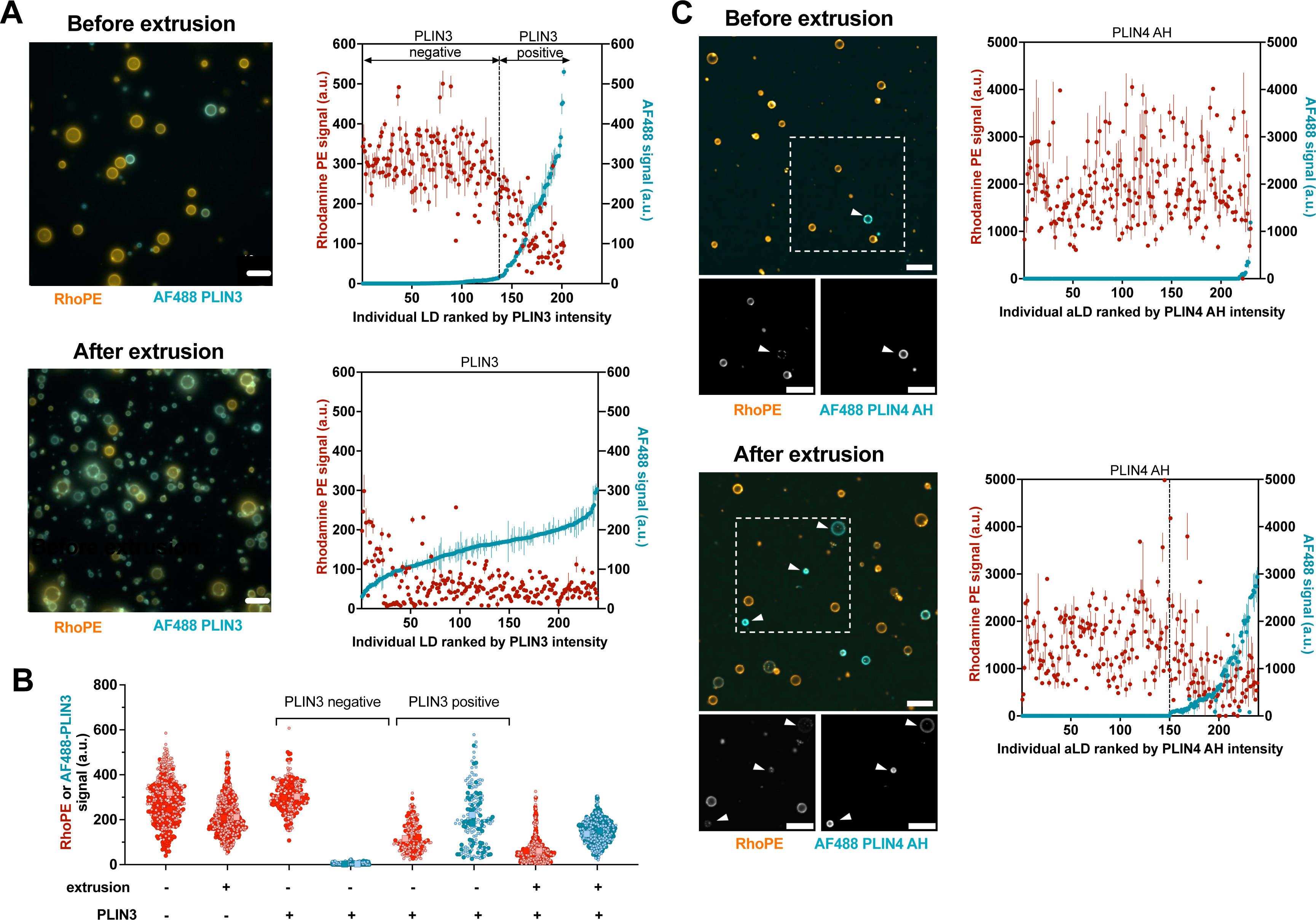
Binding of PLIN3/4 to PC-covered LDs negatively correlates with phospholipid density. **A.** Left: typical wide field fluorescence microscopy images of PC(16:0/18:1)-covered LDs (10 µm; with 0.5 % RhoPE shown in orange) in the presence of PLIN3 (with 10% AF488 PLIN3 shown in blue) before and after extrusion. Plots on the right show the individual fluorescence intensities of AF488 Plin3 and RhoPE on 200 - 250 droplets as ranked according to AF188 Plin3 intensity. **B.** LD surface analysis of the phospholipid and protein densities from two independent experiments similar to that shown in A. Data are shown as superplots (69). Light and dark symbols distinguish the two experiments. Each small circle is one LD. The large squares show the mean. **C**. Same analysis as in (A) with PLIN4 AH (12-mer) except that the images were acquired with a confocal microscope. Scale bars: 10 µm.

We performed a similar analysis with the AH region of PLIN4, for which previous studies demonstrated that it could directly coat and emulsify triolein in the absence of phospholipids and that it could rescue the size of LDs in cells after PC depletion (19, 20). PLIN4 AH (12 mer) showed an even higher sensitivity to extrusion than PLIN3. Before extrusion, only a few % of LDs showed a detectable AF488 PLIN4 AH signal. After extrusion about 25% LDs became 488 PLIN4 AH positive. Furthermore, we observed a strong anticorrelation between the AF488 PLIN4 AH and the Rho-PE signals (**Figure 4C**).

### Visualization of PLIN3 binding to bilayers does not reveal phospholipid exclusion

Given the link between PLIN3 binding to LDs and low phospholipid density, we wanted to test whether a similar effect applies when PLIN3 binds to lipid bilayers. A rare example of phospholipid exclusion in bilayers has recently been provided by the structure of caveolin, which occupies only one bilayer leaflet (46). We visualized PLIN3 on glass bead-supported lipid bilayers. In agreement with the NBD assays using liposomes (see **Figure 1D** and (15)), PLIN3 bound poorly to PC(14:0/14:0), PC(16:0/18:1) and PC(18:1/18:1)-supported bilayers, whereas strong binding was observed on supported bilayers made of PC(4Me) (**Figure 5A**). In contrast with our observations on LDs, the RhoPE signal and the AF488 PLIN3 signal coincided on all bead-supported bilayers, and we observed no significant decrease in the RhoPE signal after PLIN3 binding to PC(4Me) bilayers (**Figure 5A**). The replacement of phospholipids by PLIN3 is therefore unique to the LD monolayer.

**Figure 5.**
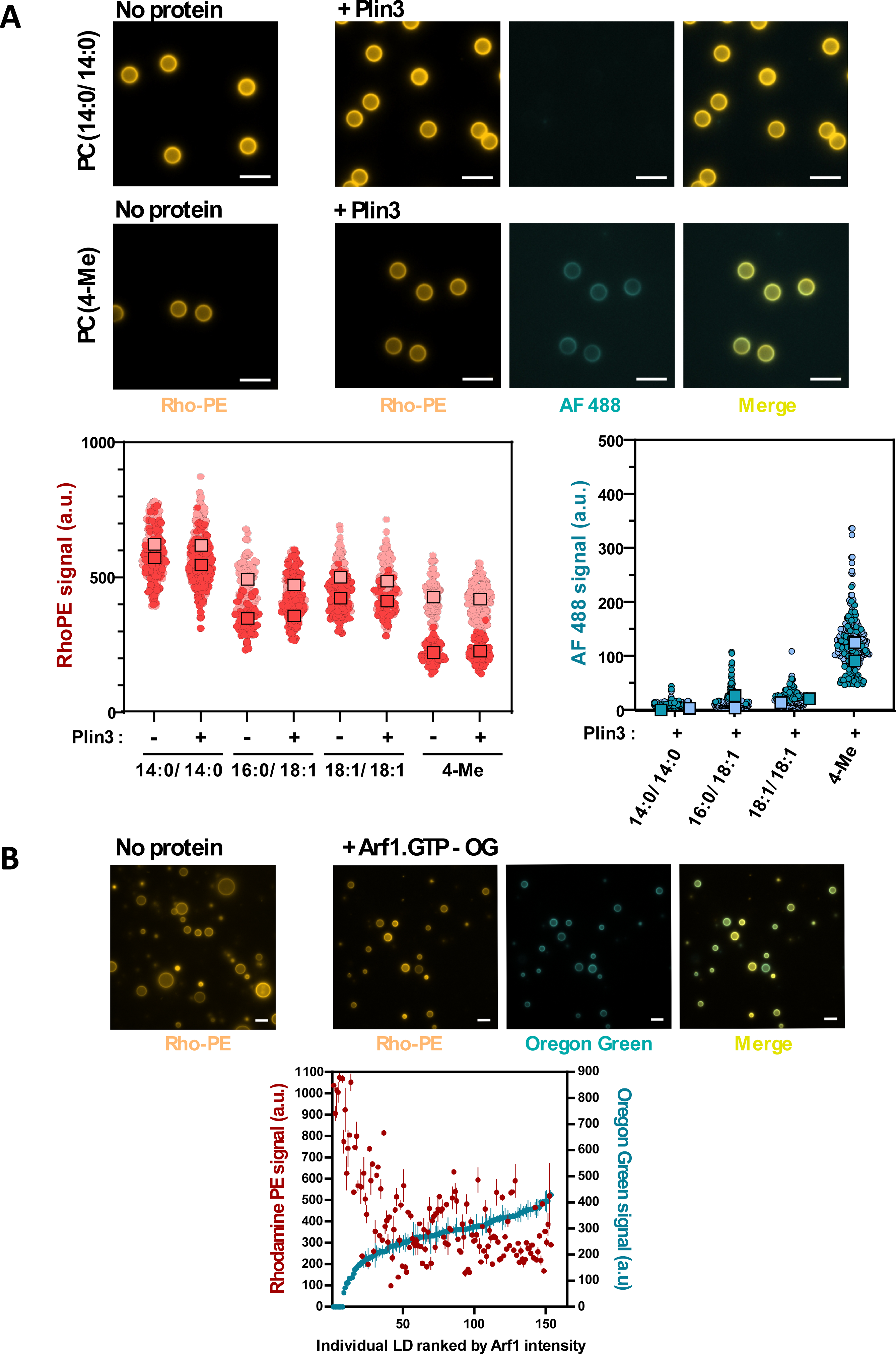
Binding of PLIN3 to PC bilayers and of Arf1 to PC-covered LDs do not correlate with change in phospholipid density. **A.** PLIN3 was incubated with 5 µm bead-supported bilayers made of PC of the indicated acyl chain composition. Images of RhoPE and AF488 PLIN3 were taken using a fluorescence microscope and quantified. Data are shown as superplots (69). Light and dark symbols distinguish two experiments. Each small circle is one template. The large squares show the mean. **B.** Arf1-OG was incubated for one hour on artificial POPC-covered LDs (containing RhoPE as a tracer) in the presence of an excess of GTPγS and in the presence of EDTA to promote nucleotide exchange. Thereafter images of Arf1-OG and Rho-PE were taken with a fluorescence microscope. The plot on the right shows the individual fluorescence intensities of Arf1-OG and RhoPE on 200 - 250 droplets as ranked according to Arf1-OG intensity.

### Visualization of Arf1-GTP binding to LDs does not reveal phospholipid exclusion

Next, we wanted to test whether other LD-binding proteins behave similarly as PLIN3 in our *in vitro* binding assay. We chose the small G protein Arf1, which has been abundantly studied at the molecular and cellular level (47). When switching to the active GTP-bound state, Arf1 exposes a myristoylated AH for organelle interaction (48). *In vitro*, Arf1-GTP binds readily to many types of lipid bilayers as well as PC-covered LDs (31). In cells, activated Arf1 can be found on many organelles, including LDs; its exact subcellular localization depends on the localization of guanine nucleotide exchange factors (49, 50). Moreover, binding of Arf1-GTP to LDs allows the recruitment of the COPI coat and the subsequent budding of nanodroplets. Such budding reduces the amounts of phospholipids at the LD surface, thereby inducing LD surface tension and facilitating the recruitment of other proteins to the LD surface (31, 50). It was thus interesting to compare the LD binding properties of Arf1-GTP and Plin3 with regard to phospholipid density.

We used an Oregon-green C-terminally labelled form of Arf1 (Arf1-OG), which binds artificial liposomes in a GTP-dependent manner akin to authentic Arf1 (51, 52). We incubated Arf1-OG with PC-covered LDs and triggered the exchange between GDP and the non-hydrolysable GTP analogue GTPγS by lowering Mg^2+^ concentration. In contrast to our observation with PLIN3 before extrusion, most LDs became intensely stained with Arf1-OG (**Figure 5B**). Variations in the Arf1-OG signal between droplets was modest and there was no obvious negative correlation between the RhoPE and the OG signals except for a minor faction of LDs that were very high in Rho-PE signal and showed almost no Arf1 signal. We conclude that binding of Arf1 to LDs is permissive to high PC monolayer density and that Arf1-GTP cannot distinguish between a monolayer and a bilayer.

Altogether, the experiments shown in Figure 5 indicate that that the replacement of phospholipids by PLIN3 at the LD surface is unique to LDs and specific proteins: neither the binding of PLIN3 to phospholipid bilayers nor the binding of Arf1-GTP to LDs leads to phospholipid exclusion.

*LD tension promotes shallow lipid packing defects that are independent of PC acyl chain profile* In lipid bilayers, replacing saturated with monounsaturated phospholipids promotes the formation of packing defects, which further increase upon membrane curvature (35). As visualized by molecular dynamics simulations, these defects appear deep, hence explaining the preferential adsorption of AH with large hydrophobic groups, such as ALPS motifs, to highly curved and monounsaturated membranes (35, 53). Interestingly, previous simulations of ternary interfaces between TG(18:1/18:1/18:1), phospholipids and water revealed that surface tension promotes the formation of different lipid packing defects from those observed in bilayers (Bacle et al., 2017; Kim and Swanson, 2020; Prévost et al., 2018). These defects are wide but shallow and result from the interdigitation of TG(18:1/18:1/18:1)within the POPC monolayer. Such defects might be more adapted to PLIN AHs, which contain rather small hydrophobic residues (e.g. Ala, Val, Thr) (20).

To extend the analysis, we performed MD simulations on ternary (TG(18:1/18:1/18:1)/PC/water) systems in which we varied both the density of the PC monolayer (surface tension) and its unsaturation level (14:0/14:0, 16:0/18:1 and 18:1/18:1) (**Figure 6A** and **S5**). At high PC coverage density (i.e. low tension), the surface of the PC monolayer was comparable to the surface of a bilayer, showing an increase in lipid packing defects according to the level of PC unsaturation (14:0/14:0 < 16:0/18:1 < 18:1/18:1). When we decreased the density of PC, the differences between the three monolayers decreased. Deep lipid packing defects increased and then plateaued, whereas shallow packing defects exponentially increased regardless of the actual unsaturation level of PC (**Figure 6B**). These defects corresponded to regions where TG(18:1/18:1/18:1) molecules were directly exposed to the solvent (6) (**Figure 6A** and **S5**). Altogether, the simulations were in line with the experimental data and reinforced the conclusion that the acyl chain profile of phospholipids is less defining for the interfacial properties of lipid droplets under tension as compared to lipid bilayers (**Figure 6C**). The LD surface packing properties depend mostly on the oil reservoir underneath, an effect that becomes prominent at low phospholipid density.

**Figure 6.**
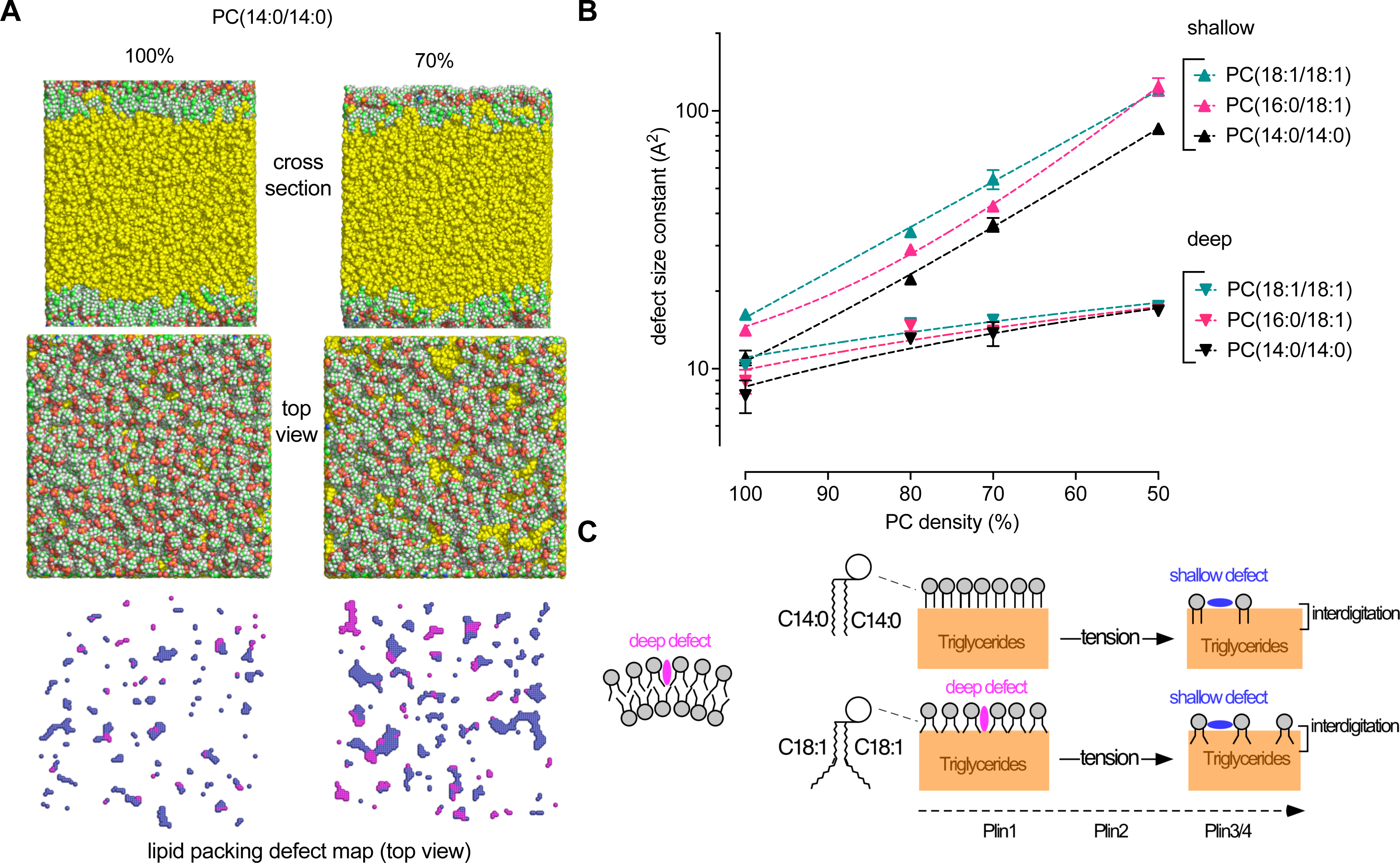
Molecular dynamic simulations of ternary water/PC/triolein systems at increasing tension. **A.** Cross section, top view and packing defect map of TG(18:1/18:1/18:1) (yellow) covered with a monolayer of PC(14:0/14:0) at 100% or 70% PC coverage (N: blue, C: green, O: red, H: white). The cartesian maps show the corresponding top view of deep (purple) and shallow (blue) defects (68). For a complete view of the combined effects of tension and PC unsaturation, see Fig. S5. **B.** Quantification of the packing defects as a function of PC coverage and unsaturation. Each point corresponds to one ternary water/PC/ TG(18:1/18:1/18:1) system with the indicated % of PC. The exponential distribution of deep and shallow defects was converted into a characteristic area constant, expressed in Å^2^ (68). **C.** Schematic view of the impact of surface tension and PC unsaturation on LD molecular surface. As surface tension increases, the underlying TG(18:1/18:1/18:1) molecules interdigitate with PC and become solvent exposed, thereby inducing the formation of shallow defects. These defects differentially condition binding of PLIN1-4 to LDs and are distinct from the deep lipid packing defects induced by curvature and mono-unsaturation on PC bilayers.

## Discussion

Although PLINS are the most abundant LD associated proteins and were among the first to be identified, our understanding of their mode of interaction with LDs has remained poorly understood. Notably, the mechanisms enabling the various PLINs to localize to different LDs in the same cell are mysterious. Here, we succeeded in purifying full-length PLIN1/2/3 and developed a novel method of LD reconstitution, which allowed us to study PLIN-LD interactions under both equilibrium conditions and following mechanical perturbations of the LD surface. Chemical and physical changes of the LD surface are inherent features of LDs and may happen at different LD life stages, from lipogenesis to lipolysis (1, 54). We also improved LD analysis by mastering LD density and by using complementary methods for quantification.

Our major conclusion is that different PLINs show very different requirements for binding to LDs. On the one hand, the interaction of PLIN1 with LDs is very robust and is affected neither by changes in LD surface composition nor by mechanical perturbations of the lipid surface, allowing PLIN1 to stably associate with LDs during adipocyte differentiation. While this manuscript was in preparation, a preprint appeared showing that PLIN1 contains two hydrophobic regions downstream of the AH that make the protein behave as an integral ER protein in the absence of LDs (55), consistent with our observations with purified PLIN1. On the other hand, PLIN3 is very sensitive to large perturbation of the LD surface; its binding requires transient LD deformation (in our *in vitro* experiments promoted by LD extrusion), an effect that largely surpasses what can be achieved by changing the lipid composition of the LD surface, including the incorporation of DOG, which has a smaller headgroup than phospholipids. PLIN2 falls between these two extreme cases. The truncated versions of PLIN4 used here suggest that, akin to PLIN3, PLIN4 is a low affinity LD protein, consistent with previous observations (19, 20). In addition, MD simulations show a poor resemblance of packing defects in model LDs to those observed on bilayers, especially at high LD surface tension when the phospholipid coverage of LDs is only partial. Instead of displaying deep cavities adapted to bulky hydrophobic residues, the LD surface at high tension shows shallow but wide lipid packing defects, which result from the interdigitation of the underneath oil molecules with the phospholipid monolayer (5, 6, 9). This interdigitation reduces the impact of the phospholipid acyl chain profile on the LD surface properties and makes LDs under tension well adapted to host extended AH protein regions containing small hydrophobic residues, a hallmark of PLIN3 and PLIN4 (1, 14, 19, 20). When we analyzed PLIN3 or PLIN4 AH on LDs, we observed a strong anti-correlation between protein and phospholipid densities, suggesting that these proteins essentially replace the phospholipid bilayer.

Interestingly, the two PLINs specific for mature adipocytes (42–44, 56) are the most different: PLIN1 is not soluble whereas PLIN4 is a gigantic intrinsically disordered soluble protein (14). In resting differentiated adipocytes, these contrasting properties translate into different localization. PLIN1 is present on the large pre-existing LDs, whereas PLIN4 is in the cytosol. However, upon fatty acid addition, the new droplets become solely decorated by PLIN4. Fast growing LDs might be better handled by a low affinity soluble PLIN because it is immediately available. In addition, PLIN4 is adapted for coating LDs with its highly extended AH (19, 20). PLIN2 and PLIN3 are more ubiquitous. Given their difference in LD affinity, it is possible that they follow the same division of labor as PLIN1 and PLIN4, with PLIN2 coating mature droplets and PLIN3 handling fast forming ones.

Overall, the use of a repertoire of PLINs with different solubility and affinity for LDs might help cells to cope with LDs of different dynamics. For example, during the process of LD formation, it has been noticed that for an LD to quickly and faithfully bud toward the cytosolic side of the ER, there must be mechanisms to control its cytosolic surface. If not, the droplet might bud toward the luminal side (57–59). In cells, impeding or boosting phospholipid metabolism can indeed affect LD size and/or budding direction, which can be corrected by expressing PLINs (19, 57). In simulations and biochemical reconstitutions, LD budding directionality can be controlled by adding new phospholipids or proteins to one bilayer leaflet in parallel with TG supply (Chorlay et al., 2019; Choudhary et al., 2018; Nieto et al., 2023). Although in such in vitro systems, strong hydrophobic proteins appear more efficient, this is tempered by limitations in solubility (57). Our study suggests that the benefit of low affinity PLINs is to provide a highly soluble pool to specifically coat LDs under tension.

The capacity to prepare artificial droplets covered with full-length perilipins should offer new possibilities to study their function, interactions and differences. These include their ability to sense different oils, the dissection of the kinase-dependent lipolysis cascade on PLIN1 LDs, or the mechanism of CIDE-induced LD fusion. The possibility to analyze lipid droplets by flow cytometry provides a level of precision that surpasses all methods since it combines the single LD level with the handling of thousands of LDs from microliter samples. The importance of understanding the precise mechanisms of PLIN function on LDs is underscored by their diverse implications in different diseases. Mutations in PLIN1 and PLIN4 in particular have been linked to many metabolic phenotypes, and loss of function heterozygous mutations in PLIN1 and PLIN4 positively and negatively, respectively, correlate with metabolic disease (14, 60). Over-expression of PLIN2 and PLIN3 has been observed in a number of cancers and often correlates with higher proliferation and poor prognosis (61). Overexpression of PLIN4 has been shown to promote drug resistance of triple-negative breast cancer (62). These links are not surprising, given the central role of LDs in mediating cellular lipid metabolism and energy homeostasis. Mechanistic understanding of PLIN function on LDs therefore has important implications for human health.

## Material and Methods

### Synthetic genes and plasmid constructions

Codon optimized sequences of human Plin1, Plin2 and Plin3 were ordered from Eurofins Genomics TM. They contained, upstream, a NdeI (CATATG) restriction site, a hexahistidine tag (CATCATCACCATCACCAC), a Tev site (GAA AACCTGTACTTCCAAAGC) and, downstream, a Ssp1 restriction site. The genes were sub-cloned into a NdeI/Ssp1 digested pET16b.His10.TEV.LIC expression vector (63). All constructs were controlled by DNA sequencing before transformation into E. coli strains.

### Protein expression and purification

The codon-optimized plasmid for hPLIN1 was expressed in *Escherichia coli* ArcticExpress (DE3) (Agilent) while codon optimized plasmids hPLIN2 and hPLIN3 were expressed in *E. coli* BL21-Gold (DE3). Cells were grown in LB medium with 50µg/mL of ampicilin at 37°C in 2L flasks to an OD_600nm_ of ≈ 0.6 and induced with 1mM isopropyl b-D-1-thiogalactopyranoside (IPTG) at 37°C for 3h in the case of hPLIN2 and hPLIN3. For hPLIN1, cells were cooled on ice for 30 min before induction with 1mM isopropyl b-D-1-thiogalactopyranoside (IPTG) at 27°C for 1h30. Cells were harvested by centrifugation for 20 min at 6000 g and stored at -20°C.

For the purification of PLIN1 and PLIN 2, pellets were resuspended in buffer A (25mM Tris, pH 7.5, 300mM NaCl, 30 mM imidazole,) containing cOmplete EDTA-free protease inhibitor cocktail (Roche), 1,5µM pestatin and 2µM bestatin. The cells were lysed using a cell disruptor, supplemented with 0.5 mM phenylmethylsulfonyl fluoride (PMSF) and centrifuged at 120 000x g at 4°C for 45min. Pellets were resuspended in Buffer B (25mM Tris-HCl, 300mM NaCl, 30mM Imidazole, 7M Urea, pH 7.5) and centrifuged at 120 000x g, at 4°C, for 30min. The resulting supernatant was incubated at 4°C for 3H with pre-equilibrated Co-NTA beads (Fisher). The unbound fraction was eluted from the beads, and the beads were washed with 10 volumes of buffer B before elution with Buffer C (20 mM Mes, 300 mM NaCl, 500 mM imidazole, 7M Urea, pH 6.3). The eluted fractions were supplemented with 2 mM dithiothreitol (DTT), pooled and concentrated on an Amicon 10kDa MWCO cell. The protein pool was purified on a sephacryl S300 HR column (Cytiva) equilibrated with Buffer D (25 mM Tris, 120 mM NaCl, 3 M Urea and 2mM DTT, pH 7.5). For 1L of culture, the average yield of purification was 0.5 and 30 mg for hPLIN1 and hPLIN2, respectively.

For the purification of hPLIN3, the bacteria pellet was resuspended in buffer A supplemented with a tablet of cOmplete EDTA-free protease inhibitor coktail (Roche), 1,5µM pestatin and 1.5 µM bestatin. Cells were lysed using a cell disruptor, supplemented with 0.5mM of PMSF, and centrifuged at 120 000x g, at 4°C, for 45min. The resulting supernatant was incubated at 4°C for 3h with pre-equilibrated Co-NTA beads (Fisher). The unbound fraction was separated from the beads, and the beads were washed with 10 volumes of buffer A before elution with Buffer E (25mM Tris, 300 mM NaCl, 500 mM imidazole, pH 7.5). The eluted fractions were supplemented with 2 mM of DTT, pooled, and incubated overnight with TEV (Tobacco Etch Virus) protease to remove the 6His-Tag. The pooled fraction was concentrated on an Amicon 10kDa MWCO cell and applied to a sephacryl S300 HR column (GE Healthcare) equilibrated with Buffer F (25mM Tris pH 7.5, 120 mM NaCl, and 2mM DTT). The average yield of full-length hPLIN3 was 7mg per liter of culture.

### Protein labelling

Labelling of PLIN3 was performed after purification. Following the gel filtration step, the protein was loaded on an Illustra NAP-5 column equilibrated with 25mM Tris, pH 7.5, 120mM NaCl. Thereafter, PLIN3 was labeled on endogenous cysteines by incubation at room temperature with a 10-fold excess of maleimide AF488 C5 or IANBD amide (N,N0-dimethyl-N- (iodoace-tyl)-N’-(7-nitrobenz-2-oxa-1,3-diazol-4-yl)ethylenediamine)) for 15min. The unconjugated probe was blocked with L-cysteine and removed by buffer exchange on a NAP-5 column.

In the case of PLIN1 and PLIN2, labelling was performed during purification. Proteins associated with Co-NTA beads were incubated at room temperature with an excess of maleimide AF488 C5 for 15min to label endogenous cysteines. The unconjugated probe was removed by washing the beads with buffer B before elution of the protein using buffer C. The labelled protein was further purified by size exclusion chromatography as described for the unlabeled form.

PLIN4 AH (12-mer or 20-mer) were purified and labelled with Alexa Fluor 488 C5 maleimide as described (19). Myristoylated Arf1 C182-Oregon green was prepared as described (51).

### Limited proteolysis

Limited proteolysis was carried out in 25 mM Tris pH7.4, 120mM NaCl, 1mM MgCl_2_ and 1 mM DTT with a final concentration of 0M or 2M Urea. Proteins (3µM) were incubated at 25°C under agitation with 1 µg/mL of subtilisin. At indicated times, aliquots were withdrawn and the reaction was stopped by adding 2 mM PMSF.

### Artificial lipid droplet preparation

All lipids were from Avanti polar lipids, except triolein (18:1/18:1/18:1) and BVO (CAS: 8016-94-2), which were from Sigma and Spectrum Chemical MFG Corp, respectively. Contaminants such as free FA were removed from BVO following a method described elsewhere (64). Briefly, polypropylene tubes were first washed with hexane for 24h. Then, 8 parts (mL) of methanol were added to 1 part of oil (g), and vortexed. The tubes were centrifuged at 1650 g for 5 min. They were then placed upright in the freezer (< -20°C) for at least 8h. The methanol was discarded and the tubes were left at RT to thaw the oil. New methanol was added and the process was repeated 5 times. Finally, the mix was dried in a rotavapor and stored at -20C in an argon enriched atmosphere.

To prepare lipid droplets of defined density (*ρ*) and diameter (*d*), we relied on the following equations:

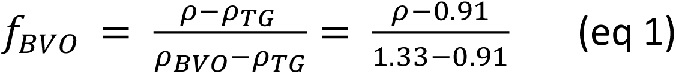

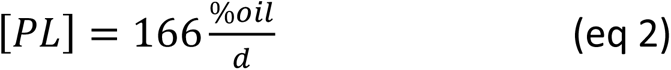

Equation 1 gives the volume fraction of BVO as a function of TG density *(π_TG_*), BVO density (*π_BVO_*), and the actual oil density (*ρ*) of the mixture. For a density *ρ* = 1.05, which was used in most experiments, this gives a volume fraction of BVO and TG of 0.33 and 0.67, respectively.

Equation 2 gives the phospholipid concentration ([*PL*], in µM) that is needed to emulsify a defined percentage of oil in buffer (*%_oil_*) into LDs with a defined average diameter (*d,* in µm). This equation derives from two calculations of the total surface of lipid droplets (*S_oil_*):

First, the total number of phospholipid molecules multiplied by their elementary surface (*A_PL_* ≈ 0.7 nm^2^) gives the surface of phospholipid monolayer available:

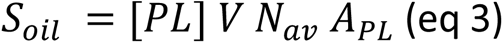

where *N_av_* is the Avogadro number (6.02 x 10^23^), *V* is the volume of the emulsion and *A_PL_* is the elementary surface of phospholipids.

Second, the total surface of lipid droplets is also the number of LDs (*n*) multiplied by their elementary surface;

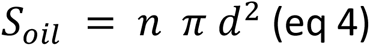

Because the number of droplets, *n,* is the ratio between the oil volume and the elementary volume of the droplets, this gives:

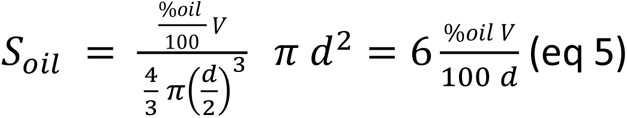

Combining equations 3 and 5 gives equation 2.

For a volume fraction of 0.75% oil in buffer, obtaining a suspension of LDs with a calculated diameter of 2 µm requires a concentration of phospholipid [PL] = 62.5 µM. This concentration was used to prepare droplets for flotation and flow cytometry measurements. For larger lipid droplets used in light microscopy experiments (calculated diameter 10 µm), we used a five-fold lower concentration: [PL] = 12.5 µM.

To prepare the LDs, we first mixed 10 µL TG(18:1/18:1/18:1) (9.11 mg) and 5 µL BVO (6.6 mg) from stock solutions in chloroform (≈ 100 mg/ml). For LDs used for flotation and flow cytometry experiments (2 µm LDs), the mix was supplemented with 125 nmol of a chosen PC species (PC(14:0/14:0), PC(16:0/18:1), or PC(18:1/18:1)) and 0.625 nmol Rhodamine-PE, both as stock solutions in chloroform. For LDs used for fluorescence microscopy experiments (10 µm LDs), the mix was supplemented with 25 nmol of a chosen PC species (PC(14:0/14:0), PC(16:0/18:1), or PC(18:1/18:1)) and 0.125 nmol Rhodamine-PE. After evaporation of chloroform under a stream of nitrogen, the final oil volume containing the phospholipids (10 µL TG and 5 µL BVO) was resuspended with 2 mL HKMD buffer (Hepes 50 mM pH 7.2, K acetate 120 mM, MgCl_2_ 1 mM, DTT 1 mM), hence leading to a 0.75% oil suspension. The suspension was vigorously vortexed for 2 minutes at maximum speed and then extruded 19 times through 1 µm (for LDs used in flotation and flow cytometry experiments) or 8 µm polycarbonate filters (for LDs used in fluorescence microscopy experiments) using a hand mini extruder (Avanti). The LDs were stored at room temperature under argon and protected from light.

### LD flotation on sucrose gradients

500 µL of 1 µm-extruded LDs (0.75 % oil) covered with the indicated PC (14:0/14:0, 16:0/18:1, or 18:1/18:1; concentration = 62.5 µM and 0.5 mol% RhoPE) were supplemented with protein (e.g. PLIN3) among which 5 - 20 % was fluorescently labelled in a total volume of 600 µL. Half of the sample was then further extruded 19 times through a 1 µm pore size polycarbonate filter. 150 µL of the non-extruded or extruded samples were then mixed with 100 µL Sucrose (75% w/v) in HKMD buffer in a centrifuge tube. The resulting 30% sucrose cushion was overlaid with 250 µL sucrose (25% w/v in HKMD buffer) and 50 µL HKMD buffer. After centrifugation for 1 hour at 55000 rpm and at 20°C in a TLS 55 (Beckman) rotor, three fractions (top, middle, bottom) were collected using a Hamilton syringe and analyzed by SDS-PAGE using direct fluorescence and or Sypro-staining and by Rhodamine fluorescence to determine protein binding to LDs and LD recovery.

### Flow cytometry experiments

Similar samples as those used in flotation and microscopy experiments were prepared and analyzed by flow cytometry. The samples containing 8 µm or 1 µm LDs were diluted 10 or 50 times, respectively, in HKMD buffer to a final volume of 500 µL. Samples were acquired with a Cytek® Aurora (Cytek Biosciences) equipped with 5 lasers (355, 405, 488, 561, 640 nm) and 64 detectors. 50 µL of each sample were recorded at medium flow rate to assess aLDs concentration and determine fluorescence intensity and percentage of LDs with bound protein. Only particles within the typical S shape area of the scattering diagrams ISS(IFS) were analyzed (see Fig. S2E). Data were unmixed in SpectroFlo v3.0.1 (Cytek Biosciences) and analyzed in FlowJo v10.9 (BD Biosciences).

We also performed similar flow cytometry experiments using an Attune CytPix flow cytometer, which is equipped with a bright field camera, to measure LD size. After gating the areas of interest, a total of 10000 images were taken per sample and were first processed in the Attune Cytometric software v6.0.1. Images with aggregated LDs were eliminated and the remaining images were analyzed in batches (per condition per gated region) using a dedicated pipeline for Cell Profiler v4.2.1 and v4.2.5. The minimum Feret diameter was chosen to avoid manual screening and elimination of images containing aggregated LDs that might have passed through the first selection filter.

### Dynamic light scattering (DLS) analysis

The same protein/artificial lipid droplet mixtures as those used for flotation or flow cytometry experiments were analyzed by DLS without dilution in a VASCO KIN particle size analyzer (Cordouan technologies).

### Lipid extraction and HPTLC

Lipids were extracted from 100 µL of small LDs, before and after extrusion, with and without protein. Because in LDs the ratio PL/TG is very low, a 3-phase liquid extraction (3PLE) was performed, in order to separate polar and neutral lipids in different organic phases. The method and solvent ratios used were adapted from (65, 66). Briefly, the separate solvents were added to the sample and then the aqueous phase (sample) was completed with water, resulting in Hex:EtAc:ACN:Aqueous (3:1:3:2). On the day of extraction, 50 µg/mL BHT was added to each solvent separately. Samples were vortexed and centrifuged at 2500 g, for 4 min, at 20°C. A fixed volume of upper phase was collected, and hexane was added (half the volume of the first extraction) to the 2 remaining phases for re-extraction. Samples were again vortexed and centrifuged, and fixed volumes of upper and middle phases were collected separately. To reduce PL loss, a re-extraction of middle phase was done with ACN and EtAc (3:1) (half the volume of first extraction). Samples were again vortexed, centrifuged and collected.

Hex (upper) phases were dried under nitrogen, diluted in 0.3 mL CHCl_3_ and 50 µL were transferred to injection vials. ACN (middle) phases were dried in a vacuum centrifuge (speedvac), and transferred to injection vials. The ACN collection tubes were rinsed twice with CHCl_3_ to avoid losing sample. The ACN vials were re-dried under nitrogen and 18 µL of CHCl_3_ were added to each.

An automatic TLC Sampler 4 (ATS4, CAMAG^®^) with a spray needle was used to apply 6 and 15 µL of Hex and ACN samples, respectively, and 3 µg standards (manually mixed, Avanti Polar Lipids^®^ and Sigma-Aldrich^®^; BVO from Spectrum Chemical^®^) onto a Merck HPTLC glass plate silica gel 60 (20×10cm, layer thickness 200 µm). The syringe was washed 3 times between samples in CHCl_3_:MeOH (50:50).

The plate was then eluted with 9 different solvent mixes. The first 5 elutions were performed in an automated multiple development chamber (AMD2, CAMAG^®^), with pre-conditioning in MeOH. The following 3 elutions were done in an automatic developing chamber (ADC2, CAMAG^®^) with a fixed humidity percentage achieved with a saturated MgCl_2_*5H_2_O solution. Saturation without pads was done for 10 min before the 2 first elutions, and plate pre-conditioning was done for 5 min. The plate was dried for 5 min between elutions. Solvent systems’ % distribution by order of elution: Ethyl acetate, 1-propanol, chloroform, methanol, 0.25% (W/V) aqueous potassium chloride, 1) 24:30:27:11:8, 2) 27:27:27:11:8, 3) 27:27:27:19:0, all up to 50 mm; 4) Ethyl acetate, chloroform (50:50), up to 55 mm; 5) Ethyl acetate, chloroform (30:70), up to 60 mm; 6) Hexane, ethyl acetate (60:40), up to 70 mm; 7) toluene up to 78 mm; 8) hexane up to 85 mm.

Plate surface was sprayed with a modified copper sulphate solution (Handloser et al., 2008) in a derivatization chamber (CAMAG^®^) and revealed by heating at 110°C in a plate heater (CAMAG^®^) under a fume hood for 40-50 min, then 5-10 min at 140°C. Imaging was done in a Fusion FX7 instrument (Vilber Loumat™), using epi white light for the charred lipids and fluorescence detection for RhoPE (before and after charring). Identification lipids was carried out by comparison to standards applied onto the same TLC plate.

Plate images were analysed in Fiji, by determining peak areas corresponding to each lipid band of interest.

### NBD fluorescence measurements with liposomes

The experiments were performed essentially as in (19). Dry films containing chosen lipids were prepared from stock solutions in chloroform. The film was resuspended in 50 mM Hepes, pH 7.2, 120 mM K-acetate at a concentration of 5 mM lipids. After five cycles of freezing in liquid nitrogen and thawing in a water bath, the multi-lamellar liposomes were extruded through 100 nm pore size polycarbonate filters. Fluorescence emission spectra upon excitation at 505 nm were recorded in a Jasco RF-8300 apparatus. The sample (600 µL) was prepared in a cylindrical quartz cell containing liposomes (0, 50, 150 or 450 µM lipids) in HK buffer supplemented with 1 mM MgCl_2_ and 1 mM DTT. The solution was stirred with a magnetic bar and the temperature of the cell holder was set at 37°C. After acquiring a blank spectrum, NBD-Plin3 (150 nM) was added and a second spectrum was measured and corrected for the blank. The fluorescence ratio of NBD-Plin3 at 540 nm in the presence of liposomes *versus* in solution was then determined.

### Protein binding to bead-supported lipid bilayers

To prepare bead-supported bilayers, extruded liposomes (200 µM lipids) were incubated with 5×10^6^ uniform silica beads of 5 µm in diameter (Bangs Laboratories, inc.) in HKMD buffer (100 µL total volume) for 30 min at room temperature under gentle mixing. The beads were washed three times in HKMD buffer and centrifuged (200×g for 2 min) before observation.

### Fluorescence microscopy of LD and bead-supported lipid bilayers

450 µL of 8 µm-extruded LDs (0.75 % oil) covered with the indicated PC (14:0/14:0, 16:0/18:1, or 18:1/18:1; concentration = 12.5 µM) were supplemented with 250 nM PLIN3 (where 5% of protein was fluorescently labelled) or PLIN4 AH (12mer, where 10% was fluorescently labelled). The total volume was 500 µL. Half of the sample was then further extruded 19 times through an 8 µm polycarbonate filter. The droplets were diluted ≈ 5 times in HKMD buffer in *Ibidi* slides previously passivated with free-FA BSA (6 mg/mL) and washed 3x with buffer. For Plin3 Images were taken with an EMCCD Camera (iXon Ultra 897, Oxford Instruments) on an inverted wide-field fluorescent microscope (IX83, Olympus) using a 60X/1.42 oil immersion objective and operated with MetaMorph (Molecular Devices). Observations of LDs with PLIN4 AH were performed with an inverted Olympus Ixplore Spin SR microscope coupled with a spinning disk CSU-W1 head (Yokogawa) using 60X UPLXAPO 1.42 NA DT 0.15 mm oil-immersion objective. Z stacks of 10 planes with a step size of 0.5 μm were acquired with an ORCA Fusion BT Digital CMOS camera (Hamamatsu) using 488 nm and 561 nm 100 mW lasers and GFP narrow (520/10) and mCherry (593/40) filters. The system was driven by CellSens Dimension 3.2 software.

For experiments with myristoylated Arf1-OG, 50 to 100 nM protein was incubated in a total volume of 200 µL in HKMD buffer in the presence of 20 to 40 µL of 8 µm-extruded LDs (0.75 % oil) and with an excess of GTPγS (40 µM). Nucleotide exchange at the surface of the LDs was promoted by the addition 2 mM EDTA, and droplets were observed in Ibidi slides as for PLIN3.

### Adipocyte Cell culture

Human adipocyte Tert-hWA cells (41) were cultured using a protocol modified from (42). Pre-adipocytes were cultured in cultured dishes on glass slides in DMEM-F12 medium (Thermo-12634010) with 10% FBS (Sigma-F7524), and supplemented with 2.5 µg/ml ßFGF (Sigma-F3685). When cells reached confluence, serum was removed from the culture medium (day P). Two days later (day 0 of differentiation) and until day 6, differentiation was initiated by the addition of an adipogenic cocktail containing 1 nM T3 (Sigma-T6397), 0.5 mM IBMX (Sigma-I5879), 5 µg/mL insuline (Sigma-I6634), 1 µM cortisol (Sigma-H0369), 1 µM dexamethason (Sigma-D4902) et 1 µM rosiglitazon (Sigma-R2408). After day 6, the adipogenic cocktail was removed and fresh culture medium with 0% FBS was added to cells every 2 days.

At different points of the differentiation (days 3, 6 and 10), cell medium was supplemented with 100 µM oleic acid (Sigma-O1383) complexed with fatty acid-free BSA (Sigma) for 2.5 hours, after which the cells were fixed and prepared for imaging.

### Preparation of protein extracts and Western blot analysis

Adipocyte cells cultured in 100 mm culture dishes were with with PBS and harvested in 300 µl of ice-cold lysis buffer (150 mM NaCl, 50 mM Tris-HCl pH 7.4, 1% NP-40, 0.5% Na-deoxycholate, 0.1% SDS, 2 mM EDTA, containing ‘Complete mix’ protease inhibitors from Roche) and passed 10 times through a 26-gauge needle. Total proteins were quantified using Bradford assay and samples were denaturated in Laemmli buffer at 95 °C for 5 min. Proteins were separated on a 12% SDS-PAGE gel and transferred to a nitrocellulose membrane (GE Healthcare), which was then inucbated over-night with primary antibodies followed by a 1-hour incubation with fluorescent secondary antibodies (Table 1), which were detected using Odyssey imaging system (Oddysey M, LI-COR).

**Table 1.** Antibodies used in this study.

| **Antibody** | **Reference** | **Dilution IF** | **Dilution WB** |
| --- | --- | --- | --- |
| **Anti-Plin1 (polyclonal Rabbit)** | Abcam-ab3526 |  | 1/1000 |
| **Anti-Plin1 (monoclonal Mouse)** | Progen-6901156S | 1/250 |  |
| **Anti-Plin2 (polyclonal Guinea Pig)** | Progen-GP41 | 1/200 | 1/250 |
| **Anti-Plin3 (polyclonal Rabbit)** | Sigma-HPA006427 | 1/250 |  |
| **Anti-Plin4 (polyclonal Rabbit)-KIAA1881** | Abcam-ab234752 | 1/250 | 1/500 |
| **Anti-HSL (polyclonal Rabbit)-D6W5S** | Cell Signaling-18381S |  | 1/500 |
| **Anti-Actin (monoclonal Mouse)** | Fisher-MA511869 |  | 1/500 |
| **Alexa anti Rabbit-488** | Fisher-A11034 | 1/500 |  |
| **Alexa anti Guinea Pig-568** | Fisher-A11075 | 1/500 |  |
| **Alexa anti mouse-647** | Fisher-A21235 | 1/500 |  |
| **Goat anti Guinea Pig, DyLight 800** | Fisher-SA5-10100 |  | 1/20000 |
| **Goat anti Mouse, DyLight 680** | Fisher-SA5-35518 |  | 1/20000 |
| **Goat anti Rabbit, DyLight 800** | Fisher-SA5-35571 |  | 1/20000 |

### Immunofluorescence and confocal microscopy

Cells were fixed in 3.2% paraformaldehyde (Thermo-28906) for 15 min at room temperature (RT). They were then gently permeabilized with 0.5% saponin in phosphate-buffered saline (PBS) containing 0.5% FBS for 15 min at RT, washed with PBS and incubated in a blocking solution containing PBS and 0.5% FBS for 30 min at RT. They were then incubated with primary antibody cocktails using antibody dilutions as indicated in Table 1 overnight at 4 °C, rinsed in PBS and incubated with secondary antibodies and DAPI stain (Fisher-D1306) for 1 hour at RT, protected from light, and mounted for imaging.

Multi-dimensional images of fixed adipocyte cells were acquired at RT with an LSM980 confocal microscope (Zeiss), using a 40X Plan-Apo 1.4NA oil-immersion objective. The microscope is equipped with T-PMT camera and driven by Zeiss Zen Blue software. Excitation sources used were: 405 nm diode laser, an Argon laser for 488 nm and 514 nm and a Helium/Neon laser for 633 nm.

### Image analysis

For artificial LDs, the images of RhoPE and AF488 channels were analyzed using a custom-made macro in Fiji. A line was manually drawn across each droplet to get both fluorescent profiles. After subtracting the background, we determined the maximal intensity of the two channels by determining the mean of the two values corresponding to the two intersections between the line and the LD contour. The analysis was repeated about 200 times for all visible droplets from 4 to 5 different fields.

Selected single z-section images of adipocytes cells were analyzed manually using Image J.

### Molecular dynamic simulations

All-atom simulations were performed using the forced field Charmm36. The triolein (TG 18:1/18:1/18:1) topology was modified as in (67). We started from bilayers (14.1 x 14.1 nm) containing 400, 412 or 440 molecules of PC(18:1/18:1), PC(16:0/18:1), or PC(14:0/14:0) in waters (50 A). We incorporated 864 molecules of triolein between the two monolayers and then performed minimization and equilibration for 110 ns. Simulations were conducted for 600 ns. Thereafter, we determined the size distribution of lipid packing defects using PackMem (68).

## ACKNOWLEDGMENTS

This study was funded by the European Research Council (ERC Synergy 856404, SPHERES) and by the Agence National de la Recherche within the project entitled Investissements d’Avenir UCAJEDI (ANR-15-IDEX-01). We thank Jacques Fattaccioli for input on flow cytometry experiments, Sophie Abelanet for help in image analysis, Bayane Sabbagh for PLIN4 12 mer purification, and Niklas Mejhert, Dominique Langin and Scott Frendo-Cumbo for comments on the manuscript. We acknowledge SABLESPlatfomes financed by the European Union through the European Regional Development Fund for the Aurora cytometer and the flow cytometry and microscopy facility from the Institut de Pharmacologie Moléculaire et Cellulaire, which is part of the « Microscopie Imagerie Cytométrie Azur» GIS IBiSA labeled platform. We also acknowledge the imaging facility MRI, member of the national infrastructure France-BioImaging supported by the French National Research Agency (ANR-10-INBS-04, «Investments for the future»).

---

**Figure S1.**
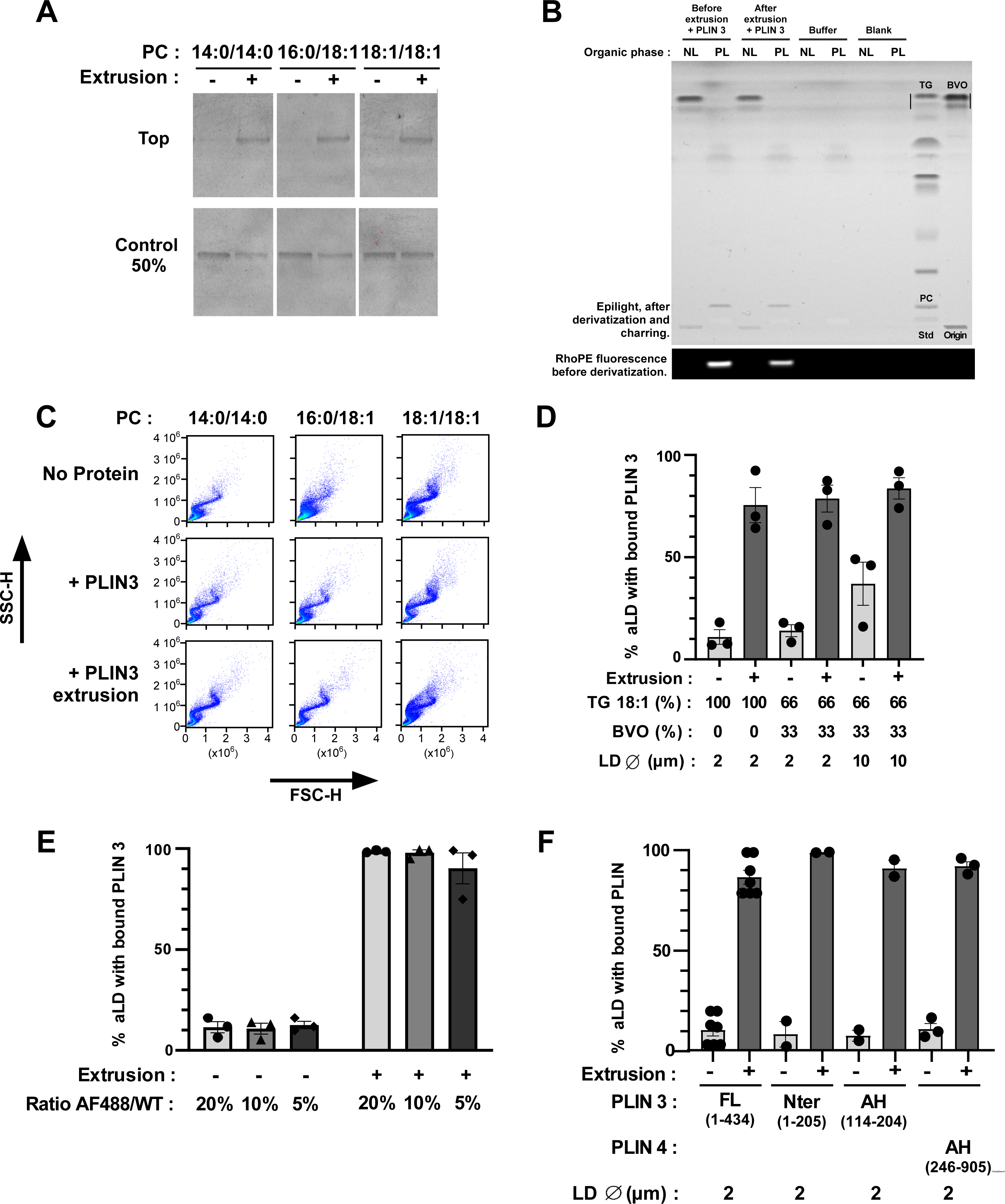
Binding of PLIN3 and the amphipathic region of PLIN3 or PLIN4 to artificial LDs requires surface tension. **A.** Effect of extrusion on the binding of PLIN3 to PC-covered lipid droplets was assessed by separating LDs from soluble protein by flotation followed by Sypro-orange staining to detect unlabeled PLIN3 in the LD-containing top fraction. Extrusion: 1 µm. **B.** HPTLC analysis of the PLIN3 + LD sample before and after extrusion (1 µm). **C**. Side scattering (SSC-H) vs forward scattering (FSC-H) diagrams of the same LDs as that shown in Figure 2C. **D.** Flow cytometry analysis of the binding of PLIN3 to PC-covered LDs made of triolein only (100%) or of triolein (66%) + BVO (34%). Droplet size: 2 µm or 10 µm. **E.** Flow cytometry analysis of PC-covered LDs in the presence of various ratio of AF488 PLIN3 *vs* unlabeled PLIN3. **F.** Flow cytometry analysis of the binding of different constructs of PLIN3 and of the AH region of PLIN4 on PC-covered LDs.

**Figure S2.**
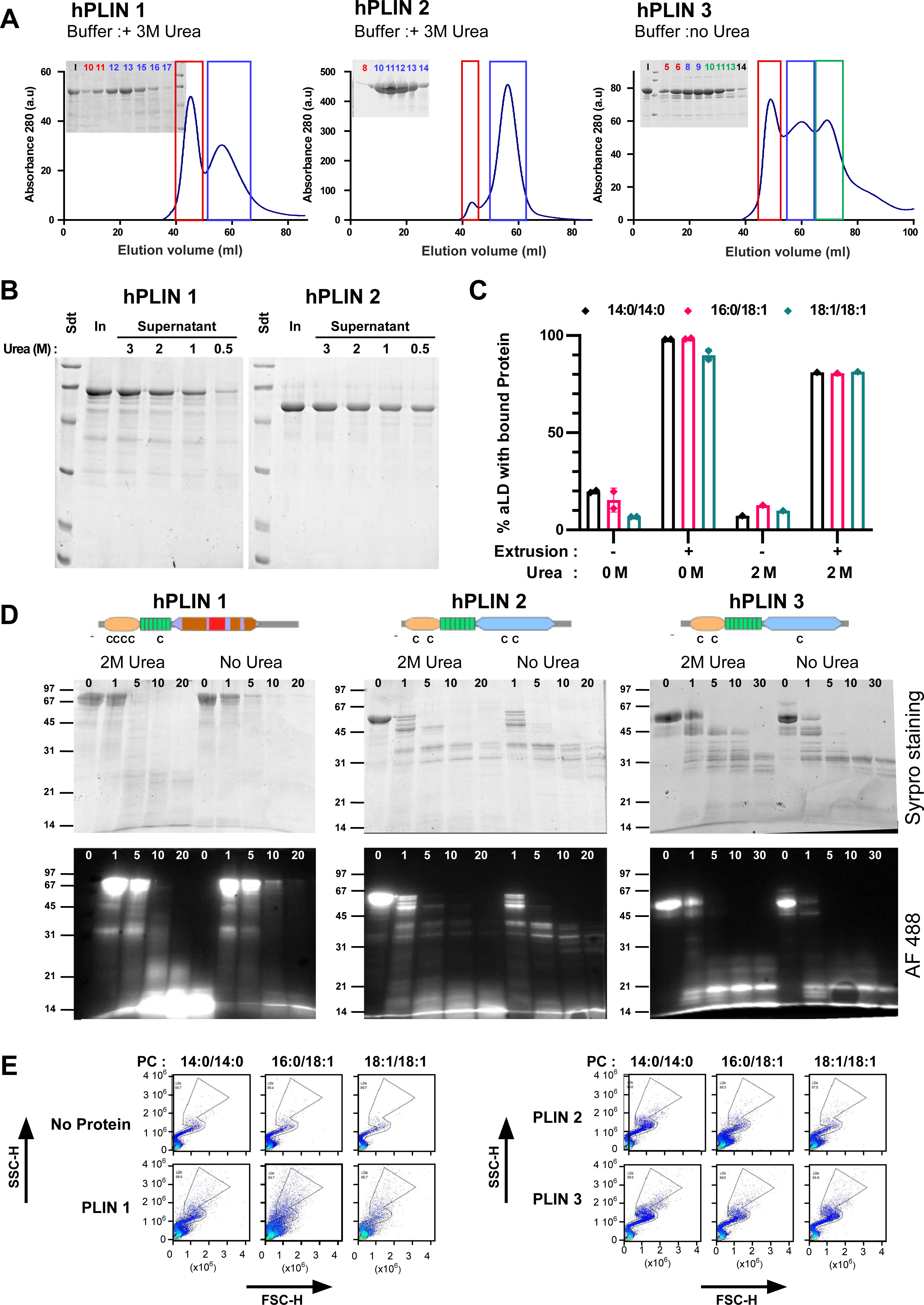
Purification and biochemical characterization of full-length human PLIN1, PLIN2 and PLIN3. **A.** Last step of purification of PLIN1-3 on a Sephacryl S300 HR column. In the case of PLIN1 and PLIN2, the buffer contained 3M urea. **B.** Sedimentation assay to determine the concentration of urea (in M) at which PLIN1 and PLIN2 remain soluble. In: input. **C**. Flow cytometry analysis of PLIN3 + PC-covered LDs in the presence or absence of 2M urea. The experimental conditions were as in Figure 1B-C. **D.** Time course of limited proteolysis of AF488 PLIN1, AF488 PLIN2, and AF488 PLIN3 in the presence of subtilisin (time in min). Proteins were analyzed either by direct fluorescence or by Sypro orange staining. The upper drawings show the predicted domain organization of PLIN1/2/3 and the localization of endogenous cysteines. **E.** Side scattering (SSC-H) vs forward scattering (FSC-H) diagrams of the same LDs as shown in Figure 2A in the absence or in the presence of the indicated PLINs. With PLIN1 and PLIN2, the scattering diagrams lost part of the typical S shape observed with oil emulsions made of individual droplets (36), suggesting partial aggregation of the particles.

**Figure S3.**
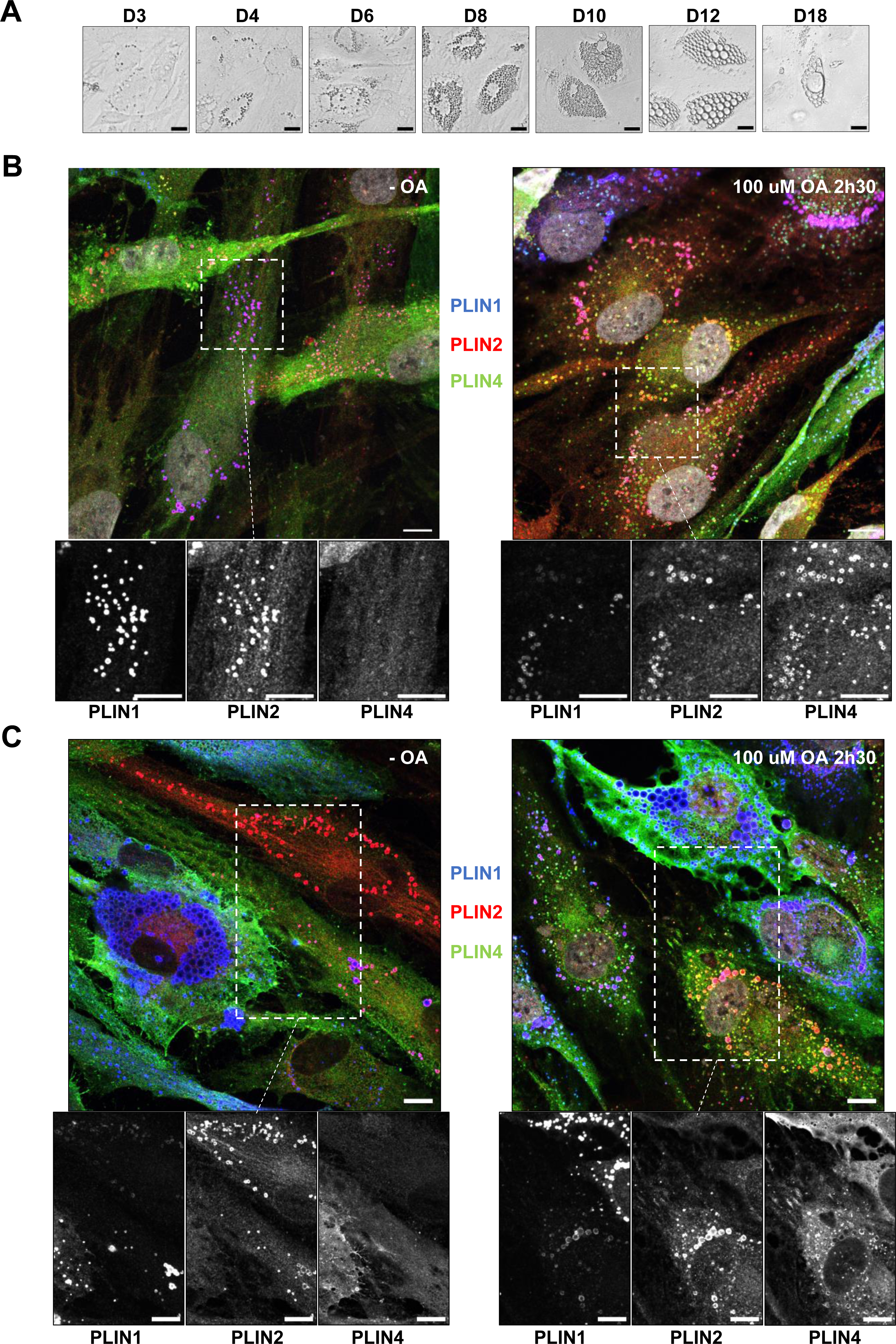
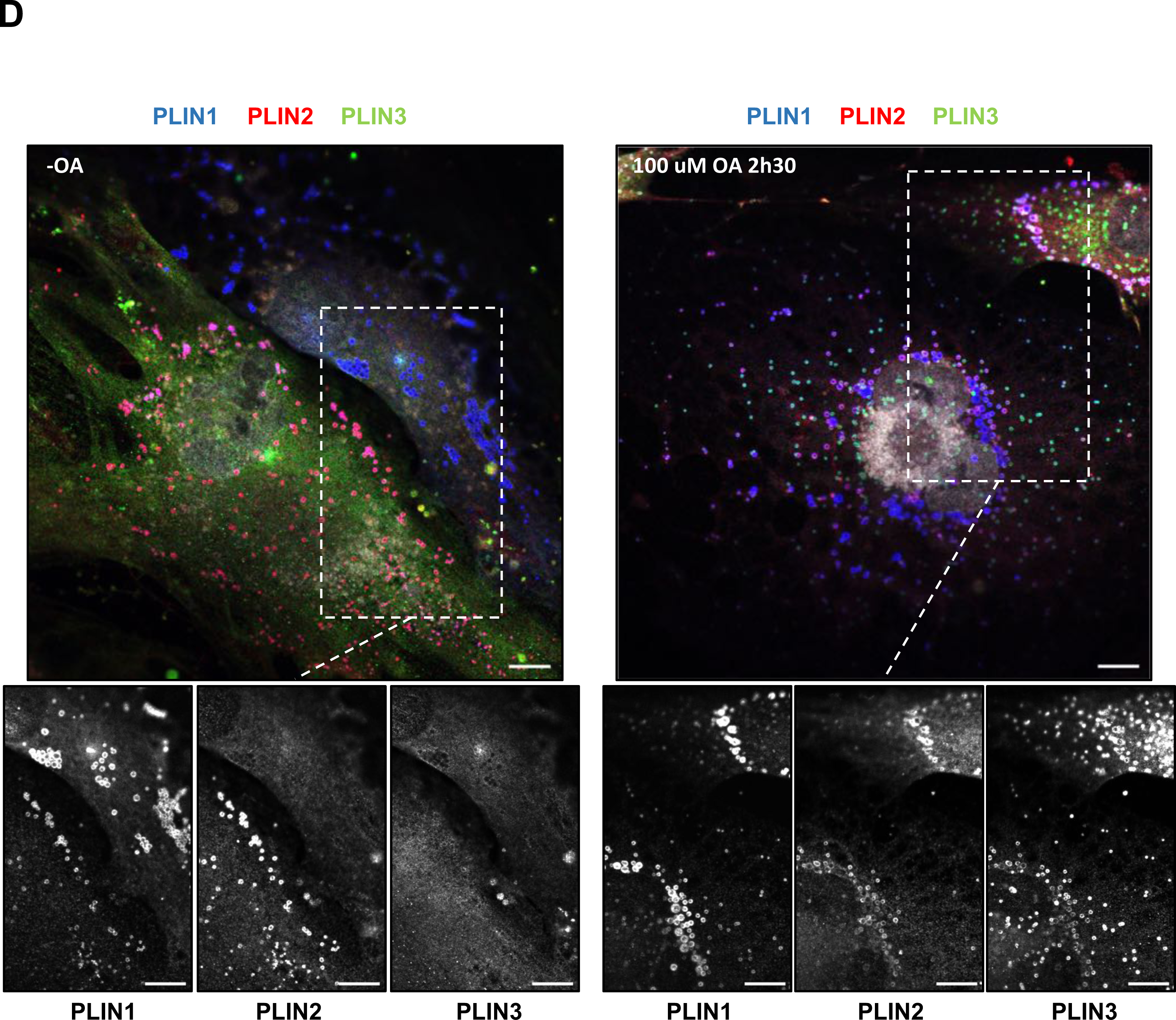
Immunofluorescence analysis of perilipins in human adipocytes during differentiation and effect of oleic acid treatment. **A.** Time course of adipocyte differentiation. **B, C.** Representative z sections obtained by confocal fluorescence microscopy of endogenous PLIN1, PLIN2 and PLIN4 in human adipocytes at day 3 (B) and 6 (C) of differentiation after immunofluorescence with specific antibodies. The cells were either left in culture medium or fed with 100 µM oleic acid for 2.5 hours before observation. Scale bar: 10 µm. **D.** same as in B and C for endogenous PLIN1, PLIN2 and PLIN3 in human adipocytes at day 6 of differentiation. Scale bar: 10 µm.

**Figure S4.**
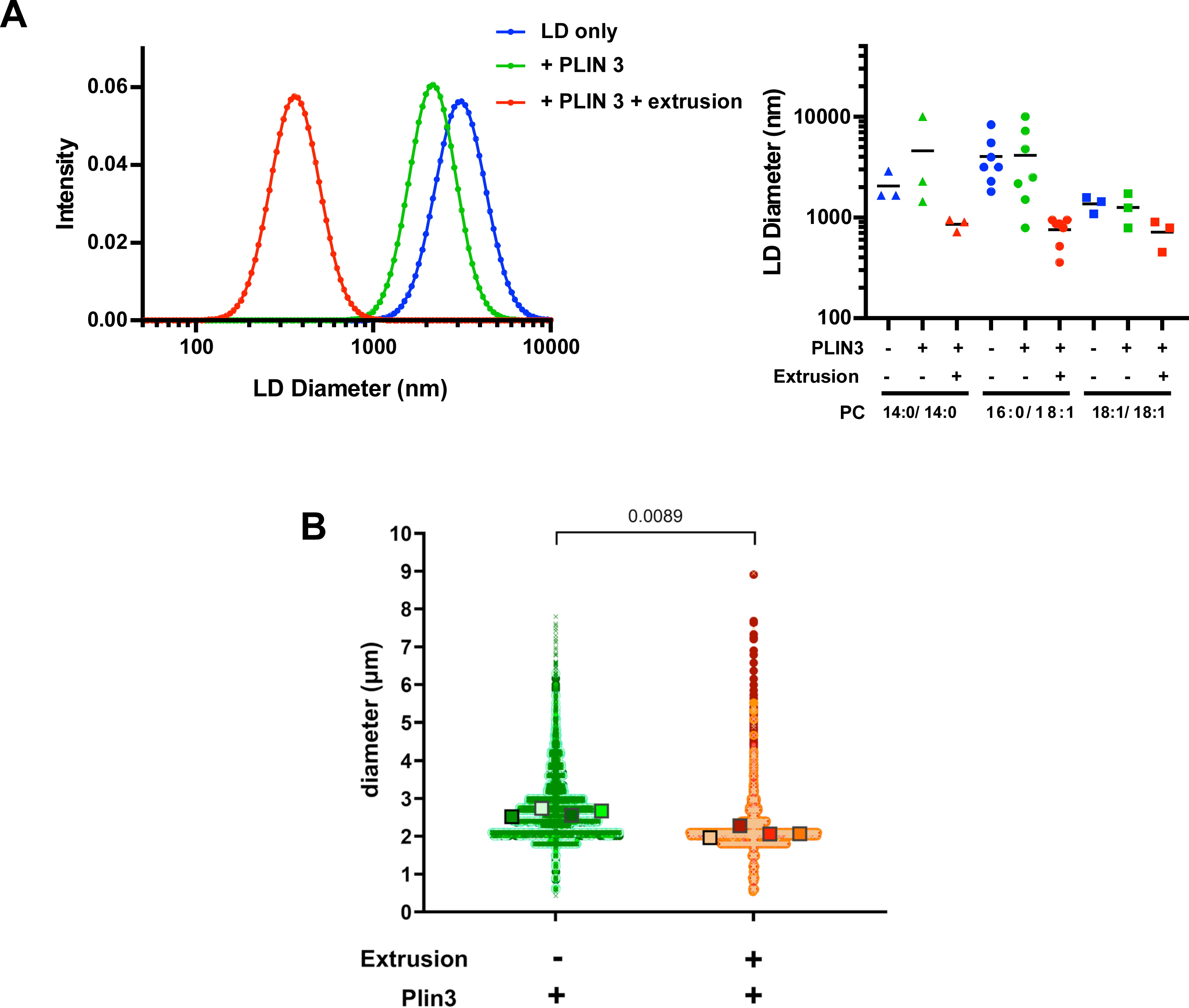
LD size analysis. **A.** DLS analysis of PC-covered LDs (calculated diameter 2 µm) in the absence or in the presence of PLIN3 and before and after extrusion (1 µm). The experimental conditions were similar as in Figure 1B and C. The left panel shows typical size distribution in one experiment using PC(16:0/18:1)-covered LDs. The right panel shows the mean diameter as determined from 3 to 7 independent experiments similar to that shown in **A. B.** Size analysis using a flow cytometer equipped with a bright field camera to measure LD size. PC(16:0/18:1)-covered LDs (calculated diameter 2 µm) were incubated with PLIN3 and eventually extruded (1 µm). After gating, the AF488 minus region was compared to the AF488 plus region, before and after extrusion, respectively, which correspond to the regions where most LDs are found (see e.g. Fig 1C). Data are shown as superplots (69). Each small circle is one droplet. The large squares show the means from four independent experiments.

**Figure S5.**
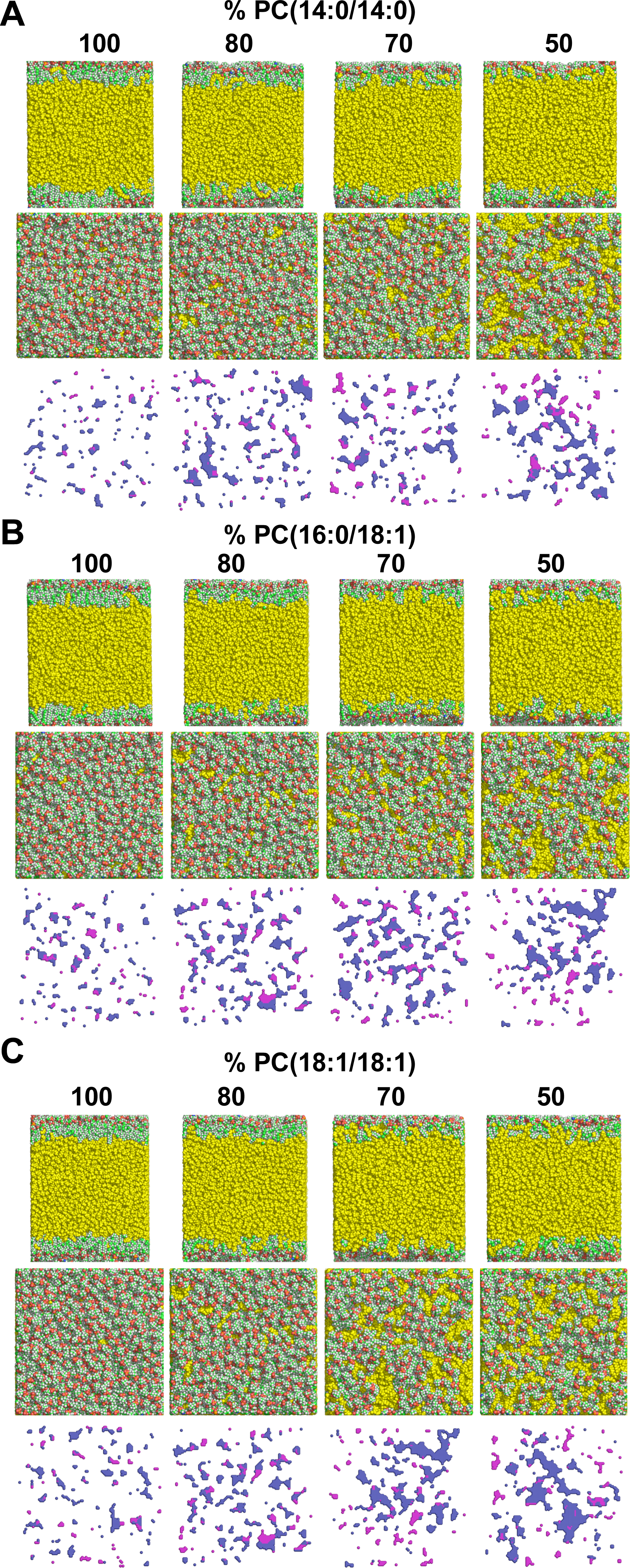
Molecular dynamic simulations of ternary water/PC/triolein systems at increasing tension. Top and side views of TG(18:1/18:1/18:1) covered with a monolayer of PC(14:0/14:0) (**A**), PC(16:0/18:1) (**B**), or PC(18:1/18:1) (**C**) at decreasing PC density. The cartesian maps show the corresponding top view of deep (purple) and shallow (blue) defects. Note the interdigitation between triolein (yellow) and phospholipids at low PC density. The difference in lipid packing defects between saturated and monounsaturated monolayers vanishes as PC density decreases.

## References

1. Dhiman R, Caesar S, Thiam AR, Schrul B (2020) Mechanisms of protein targeting to lipid droplets: A unified cell biological and biophysical perspective. Semin Cell Dev Biol 108:4–13.

2. Thiam AR, Farese RV, Walther TC (2013) The biophysics and cell biology of lipid droplets. Nature Reviews Molecular Cell Biology 14(12):775–786.

3. Chitraju C, et al. (2012) Lipidomic analysis of lipid droplets from murine hepatocytes reveals distinct signatures for nutritional stress. J Lipid Res 53(10):2141–2152.

4. Bartz R, et al. (2007) Lipidomics reveals that adiposomes store ether lipids and mediate phospholipid traffic . The Journal of Lipid Research 48(4):837–847.

5. Bacle A, Gautier R, Jackson CL, Fuchs PFJ, Vanni S (2017) Interdigitation between Triglycerides and Lipids Modulates Surface Properties of Lipid Droplets. Biophys J 112(7):1417–1430.

6. Kim S, Swanson JMJ (2020) The Surface and Hydration Properties of Lipid Droplets. Biophys J 119(10):1958–1969.

7. Bigay J, Antonny B (2012) Curvature, lipid packing, and electrostatics of membrane organelles: defining cellular territories in determining specificity. Developmental Cell 23(5):886–895.

8. Vamparys L, et al. (2013) Conical Lipids in Flat Bilayers Induce Packing Defects Similar to that Induced by Positive Curvature. Biophysj 104(3):585–593.

9. Prévost C, et al. (2018) Mechanism and Determinants of Amphipathic Helix-Containing Protein Targeting to Lipid Droplets. Developmental Cell 44(1):73–86.e4.

10. Brasaemle DL, Dolios G, Shapiro L, Wang R (2004) Proteomic analysis of proteins associated with lipid droplets of basal and lipolytically stimulated 3T3-L1 adipocytes. Journal of Biological Chemistry 279(45):46835–46842.

11. Xu S, et al. (2019) Perilipin 2 and lipid droplets provide reciprocal stabilization. Biophysics Reports 5(3):145–160.

12. Sztalryd C, Brasaemle DL (2017) The perilipin family of lipid droplet proteins_ Gatekeepers of intracellular lipolysis. BBA - Molecular and Cell Biology of Lipids 1862(Part B):1221–1232.

13. Najt CP, Devarajan M, Mashek DG (2022) Perilipins at a glance. Journal of Cell Science 135(5). doi:10.1242/jcs.259501.

14. Griseti E, et al. (2023) Molecular mechanisms of perilipin protein function in lipid droplet metabolism. FEBS Letters. doi:10.1002/1873-3468.14792.

15. Choi YM, et al. (2023) Structural insights into perilipin 3 membrane association in response to diacylglycerol accumulation. Nature Communications 14(1):3204.

16. Bussell R, Eliezer D (2003) A Structural and Functional Role for 11-mer Repeats in α- Synuclein and Other Exchangeable Lipid Binding Proteins. Journal of Molecular Biology 329(4):763–778.

17. Bulankina AV, et al. (2009) TIP47 functions in the biogenesis of lipid droplets. The Journal of Cell Biology 185(4):641–655.

18. Rowe ER, et al. (2016) Conserved Amphipathic Helices Mediate Lipid Droplet Targeting of Perilipins 1-3. J Biol Chem 291(13):6664–6678.

19. Čopič A, et al. (2018) A giant amphipathic helix from a perilipin that is adapted for coating lipid droplets. Nature Communications 9(1):1332.

20. Giménez-Andrés M, et al. (2021) Exceptional stability of a perilipin on lipid droplets depends on its polar residues, suggesting multimeric assembly. eLife 10. doi:10.7554/eLife.61401.

21. Hickenbottom SJ, Kimmel AR, Londos C, Hurley JH (2004) Structure of a Lipid Droplet Protein. Structure 12(7):1199–1207.

22. Subramanian V, Garcia A, Sekowski A, Brasaemle DL (2004) Hydrophobic sequences target and anchor perilipin A to lipid droplets. The Journal of Lipid Research 45(11):1983–1991.

23. Granneman JG, Moore H-PH, Krishnamoorthy R, Rathod M (2009) Perilipin controls lipolysis by regulating the interactions of AB-hydrolase containing 5 (Abhd5) and adipose triglyceride lipase (Atgl). J Biol Chem 284(50):34538–34544.

24. Gandotra S, et al. (2011) Human frame shift mutations affecting the carboxyl terminus of perilipin increase lipolysis by failing to sequester the adipose triglyceride lipase (ATGL) coactivator AB-hydrolase-containing 5 (ABHD5). J Biol Chem 286(40):34998– 35006.

25. Ruggieri A, et al. (2020) Multiomic elucidation of a coding 99-mer repeat-expansion skeletal muscle disease. Acta Neuropathol 590:4171–5.

26. Wang H, et al. (2011) Perilipin 5, a lipid droplet-associated protein, provides physical and metabolic linkage to mitochondria. J Lipid Res 52(12):2159–2168.

27. Ajjaji D, et al. (2019) Dual binding motifs underpin the hierarchical association of perilipins1-3 with lipid droplets. Mol Biol Cell 30(5):703–716.

28. Najt CP, et al. (2014) Structural and functional assessment of perilipin 2 lipid binding domain(s). Biochemistry 53(45):7051–7066.

29. Sincock PM, et al. (2003) Self-assembly is important for TIP47 function in mannose 6-phosphate receptor transport. Traffic 4(1):18–25.

30. Wang Y, et al. (2016) Construction of Nanodroplet/Adiposome and Artificial Lipid Droplets. ACS Nano 10(3):3312–3322.

31. Thiam AR, et al. (2013) COPI buds 60-nm lipid droplets from reconstituted water-phospholipid-triacylglyceride interfaces, suggesting a tension clamp function. Proc Natl Acad Sci USA 110(33):13244–13249.

32. Titus AR, et al. (2021) The C-Terminus of Perilipin 3 Shows Distinct Lipid Binding at Phospholipid-Oil-Aqueous Interfaces. Membranes (Basel) 11(4):265.

33. Julien JA, Pellett AL, Shah SS, Wittenberg NJ, Glover KJ (2021) Preparation and characterization of neutrally-buoyant oleosin-rich synthetic lipid droplets. Biochim Biophys Acta Biomembr 1863(8):183624.

34. Hynson RMG, Jeffries CM, Trewhella J, Cocklin S (2012) Solution structure studies of monomeric human TIP47/perilipin-3 reveal a highly extended conformation. Proteins 80(8):2046–2055.

35. Vanni S, Hirose H, Barelli H, Antonny B, Gautier R (2014) A sub-nanometre view of how membrane curvature and composition modulate lipid packing and protein recruitment. Nature Communications 5:4916.

36. Fattaccioli J, et al. (2009) Size and fluorescence measurements of individual droplets by flow cytometry. Soft Matter 5:2232–2238.

37. Skinner JR, et al. (2009) Diacylglycerol enrichment of endoplasmic reticulum or lipid droplets recruits perilipin 3/TIP47 during lipid storage and mobilization. J Biol Chem 284(45):30941–30948.

38. Ben M’barek K, et al. (2017) ER Membrane Phospholipids and Surface Tension Control Cellular Lipid Droplet Formation. Developmental Cell 41(6):591–604.e7.

39. Stribny J, Schneiter R (2023) Binding of Perilipin 3 to membranes containing diacylglycerol is mediated by conserved residues within its PAT domain. J Biol Chem:105384.

40. Garten M, et al. (2015) Methyl-branched lipids promote the membrane adsorption of α-synuclein by enhancing shallow lipid-packing defects. Phys Chem Chem Phys 17(24):15589–15597.

41. Markussen LK, et al. (2017) Characterization of immortalized human brown and white pre-adipocyte cell models from a single donor. PLoS ONE 12(9):e0185624.

42. Klingelhuber F, et al. (2023) A Spatiotemporal Proteomic Map of Human Adipogenesis. bioRxiv:2023.07.01.547321.

43. Wolins NE, et al. (2003) Adipocyte protein S3-12 coats nascent lipid droplets. Journal of Biological Chemistry 278(39):37713–37721.

44. Barneda D, Christian M (2017) Lipid droplet growth: regulation of a dynamic organelle. Current Opinion in Cell Biology 47:9–15.

45. Wolins NE, et al. (2005) S3-12, Adipophilin, and TIP47 package lipid in adipocytes. Journal of Biological Chemistry 280(19):19146–19155.

46. Porta JC, et al. (2022) Molecular architecture of the human caveolin-1 complex. Sci Adv 8(19):eabn7232.

47. Donaldson JG, Jackson CL (2011) ARF family G proteins and their regulators: roles in membrane transport, development and disease. Nature Reviews Molecular Cell Biology 12(6):362–375.

48. Liu Y, Kahn RA, Prestegard JH (2010) Dynamic structure of membrane-anchored Arf*GTP. Nat Struct Mol Biol 17(7):876–881.

49. Bouvet S, Golinelli-Cohen M-P, Contremoulins V, Jackson CL (2013) Targeting of the Arf-GEF GBF1 to lipid droplets and Golgi membranes. Journal of Cell Science 126(Pt 20):4794–4805.

50. Wilfling F, et al. (2014) Arf1/COPI machinery acts directly on lipid droplets and enables their connection to the ER for protein targeting. eLife 3:e01607.

51. Manneville J-B, et al. (2008) COPI coat assembly occurs on liquid-disordered domains and the associated membrane deformations are limited by membrane tension. Proceedings of the National Academy of Sciences 105(44):16946–16951.

52. Ambroggio EE, et al. (2010) ArfGAP1 generates an Arf1 gradient on continuous lipid membranes displaying flat and curved regions. The EMBO Journal 29(2):292–303.

53. Bigay J, Casella J-F, Drin G, Mesmin B, Antonny B (2005) ArfGAP1 responds to membrane curvature through the folding of a lipid packing sensor motif. The EMBO Journal 24(13):2244–2253.

54. Walther TC, Kim S, Arlt H, Voth GA, Farese RV (2023) Structure and function of lipid droplet assembly complexes. Curr Opin Struct Biol 80:102606.

55. Majchrzak M, et al. (2023) Perilipin membrane-association determines lipid droplet heterogeneity in differentiating adipocytes. bioRxiv:2023.10.30.564726.

56. Bäckdahl J, et al. (2021) Spatial mapping reveals human adipocyte subpopulations with distinct sensitivities to insulin. Cell Metabolism 33(9):1869–1882.e6.

57. Chorlay A, et al. (2019) Membrane Asymmetry Imposes Directionality on Lipid Droplet Emergence from the ER. Developmental Cell 50(1):25–42.e7.

58. Choudhary V, et al. (2018) Architecture of Lipid Droplets in Endoplasmic Reticulum Is Determined by Phospholipid Intrinsic Curvature. Current Biology 28(6):915–925.e9.

59. Nieto V, Crowley J, Santos D, Monticelli L (2023) Birth of an organelle: molecular mechanism of lipid droplet biogenesis. bioRxiv:2023.07.28.550987.

60. Duan X, Savage DB (2023) The role of lipid droplet associated proteins in inherited human disorders. FEBS Letters. doi:10.1002/1873-3468.14779.

61. Safi R, Menéndez P, Pol A (2024) Lipid droplets provide metabolic flexibility for cancer progression. FEBS Letters. doi:10.1002/1873-3468.14820.

62. Sirois I, et al. (2019) A Unique Morphological Phenotype in Chemoresistant Triple-Negative Breast Cancer Reveals Metabolic Reprogramming and PLIN4 Expression as a Molecular Vulnerability. Mol Cancer Res 17(12):2492–2507.

63. Jamecna D, et al. (2019) An Intrinsically Disordered Region in OSBP Acts as an Entropic Barrier to Control Protein Dynamics and Orientation at Membrane Contact Sites. Developmental Cell. doi:10.1016/j.devcel.2019.02.021.

64. Lebo JA, et al. (2004) Purification of triolein for use in semipermeable membrane devices (SPMDs). Chemosphere 54(8):1217–1224.

65. Vale G, et al. (2019) Three-phase liquid extraction: a simple and fast method for lipidomic workflows. J Lipid Res 60(3):694–706.

66. Shibusawa Y, et al. (2006) Three-phase solvent systems for comprehensive separation of a wide variety of compounds by high-speed counter-current chromatography. Journal of Chromatography A 1133(1-2):119–125.

67. Campomanes P, Prabhu J, Zoni V, Vanni S (2021) Recharging your fats: CHARMM36 parameters for neutral lipids triacylglycerol and diacylglycerol. Biophys Rep (N Y) 1(2):None.

68. Gautier R, et al. (2018) PackMem: A Versatile Tool to Compute and Visualize Interfacial Packing Defects in Lipid Bilayers. Biophys J 115(3):436–444.

69. Lord SJ, Velle KB, Mullins RD, Fritz-Laylin LK (2020) SuperPlots: Communicating reproducibility and variability in cell biology. Journal of Cell Biology 219(6):94–10.

